# A positive feedback loop between sensory and octopaminergic neurons underlies nociceptive plasticity in *Drosophila* larvae

**DOI:** 10.1101/2025.07.17.665309

**Authors:** Jean-Christophe Boivin, Yi Q. Zhao, Jiayi Zhu, Jared T. Dakin, Jing Ning, Tomoko Ohyama

## Abstract

Adaptive modulation of nociceptive behaviour based on prior experience is essential for responding effectively to environmental threats. In *Drosophila* larvae, nociceptive escape behaviours are robust and stereotyped, yet emerging evidence suggests this can be modulated by experience and internal state. Here, we demonstrate that repeated activation of nociceptive sensory neurons enhances both the likelihood and intensity of nocifensive rolling, reflecting a form of behavioural sensitization. This heightened responsiveness is accompanied by a sustained increase in activity within nociceptive sensory neurons, suggesting that plasticity arises, at least in part, within the sensory compartment. We identified the neuromodulator octopamine as a critical regulator of the sensitization: signalling through the octopamine receptor OAMB is required to sustain elevated nociceptive gain, and feedback from one of octopaminergic neurons class, the ventral unpaired median (VUM) neurons, amplifies sensory neuron output. Together, these findings reveal an experience-dependent positive feedback loop in the nociceptive system, where neuromodulatory circuits tune behavioural output.

## Introduction

The ability to modify behaviour based on experience is essential for survival in complex and ever-changing environments. This behavioural flexibility arises from the intricate interplay between an organism’s genetically determined neural architecture (“nature”) and the influence of external conditions and past experiences (“nurture”). Unraveling how these factors interact to produce adaptive behavioural outcomes remains a central goal in neuroscience, requiring a deep understanding of the molecular, cellular, and circuit-level processes that mediate experience-dependent change.

Decades of research have explored the biological mechanisms that support experience-dependent behavioural change, with synaptic plasticity emerging as a central process—particularly in the formation of associations between stimuli or events—across both vertebrate and invertebrate model systems^1–3^. In parallel, technological advances such as high-resolution electron microscopy and optogenetics have enabled detailed mapping and manipulation of entire neural circuits in small brains^4–6^. These tools have opened new opportunities to investigate how specific patterns of connectivity and activity give rise to behavioural outcomes^4–6^. Notably, such studies have highlighted the importance of neuromodulators—chemical signals released more broadly and over longer timescales than classical synaptic neurotransmitters—in dynamically regulating circuit function and enhancing behavioural adaptability^7–9^.

Neuromodulators such as dopamine, serotonin, and various neuropeptides play a pivotal role in shaping the functional architecture of neural circuits. In contrast to fast synaptic transmission, neuromodulatory signals exert temporally and spatially diffuse effects, modulating synaptic strength, circuit excitability, and overall behavioural state^7,10,11^. These modulators are uniquely positioned to act as molecular bridges between an organism’s genetic blueprint and its experiential history, biasing neural computations in a context-dependent manner^12,13^. Despite increasing recognition of their significance, the precise mechanisms through which neuromodulators drive experience-dependent behavioural plasticity remain incompletely understood.

Invertebrate model systems offer powerful platforms for dissecting these pathways, particularly due to their relatively simple and well-characterized nervous systems. In *Drosophila melanogaster*, the biogenic amine octopamine serves as a major neuromodulator and is widely considered the functional analogue of noradrenaline in mammals. Octopamine modulates a broad spectrum of physiological and behavioural processes across developmental stages, including locomotion, reproduction, aggression, sleep, metabolism, and various forms of behavioural plasticity^14–20^. Its effects are mediated through a suite of G-protein-coupled receptors (GPCRs), including OAMB, Octα2R, and three OctβRs, which activate distinct intracellular signaling cascades and influence diverse neural substrates^21^. By targeting specific circuits in a context-sensitive manner, octopaminergic signaling plays a central role in aligning behavioural output with environmental demands and internal state^19,20,22,23^.

Building on this foundation, the *Drosophila melanogaster* larval system offers a valuable context for examining how neuromodulatory influences shape behavioural responses to ecologically relevant challenges. As a species that occupies a wide range of geographic regions, *Drosophila* larvae are routinely exposed to diverse and often adverse environmental conditions^24^. During development, they encounter various noxious stimuli—including chemical irritants, extreme temperatures, and mechanical injury from parasitic wasps—that threaten their survival^25–29^. To cope with such threats, larvae rely on a well-characterized nociceptive system, in which class IV dendritic arborization (C4da) sensory neurons detect damaging stimuli and drive stereotyped escape behaviours through downstream motor circuits^28–40^. While these responses are robust, accumulating evidence suggests they are not hardwired reflexes; rather, they exhibit substantial plasticity shaped by prior experience. This modulation, which includes both sensitization and habituation, is influenced by factors such as the developmental stage of the larva^41,42^, the type of noxious input^35,43,44^, and critically, the neuromodulatory state of the nociceptive network^33,37,43,45–48^. However, the interplay between these variables remains poorly understood. Gaining a comprehensive view of how intrinsic and extrinsic factors converge to shape plasticity in nociceptive behaviour is essential for understanding how adaptive or maladaptive responses to environmental threats are sculpted over time—particularly under conditions of repeated or unpredictable stimulation^49–52^.

Here, we investigate how nociceptive experience dictates both the magnitude and directionality of nociceptive adaptation. We show that increased nocifensive behaviour correlates with enhanced sensitivity to noxious stimuli, which in turn is sustained by elevated activity within nociceptive sensory neurons. Using RNA interference, we demonstrate that the octopamine receptor OAMB is required for experience-dependent sensitization. Moreover, we find that octopaminergic feedback to nociceptive sensory neurons is both necessary and sufficient for this form of plasticity. Specifically, octopaminergic ventral unpaired median neurons (VUMs) are critical regulators of this feedback loop. Together, our findings reveal that experience-dependent sensitization is driven by an octopaminergic positive feedback mechanism between sensory neurons and their downstream modulatory partners

## Results

### Previous noxious experiences modulate responses to noxious stimuli

A previous study in *D. melanogaster* larvae reported that continuous noxious chemical stimulation of class IV dendritic arborization (C4da) neurons results in behavioural desensitization^43^. To investigate how different stimulation patterns might influence the larval nociceptive system and behaviour, we employed optogenetic activation of C4da neurons during development. Larvae expressing CsChrimson in C4da neurons were exposed to 620-nm LED (5.6 mW/mm^2^) for five seconds every five minutes from the embryo stage until the late third instar stage (Fig. 1A). We then assessed nocifensive behaviour by optogenetic activating C4da neurons and quantifying individual larval responses–specially rolling–using machine-learning-based software^53,54^ (Fig. 1B). Larvae that were developmentally simulated for 120 hours exhibited a markedly enhanced nocifensive rolling compared naive controls (Fig. 1B). While only 42% of naïve larvae responded with rolling, 98% of developmentally stimulated larvae displayed rolling behaviour (Fig. 1C). Furthermore, the latency to initiate rolling was significantly reduced in the stimulated group, with the 3.63 ± 0.52 seconds earlier than in naïve larvae (stimulated: 0.67 ± 0.03 s; naïve: 4.30 ± 0.49) (Fig. 1D). Once initiated, rolling persisted for a significantly longer duration in stimulated larvae, who spend an average of 10.11 ± 0.31 s engaged in rolling during the 30-second stimulation period, compared to 2.73 ± 0.30 s in naïve animals (Fig. 1E). These findings indicate that repeated nociceptive stimulation of C4da neurons during development induces behavioural sensitization, leading to heightened response and prolonged engagement in nocifensive behaviour.

**Figure 1.**
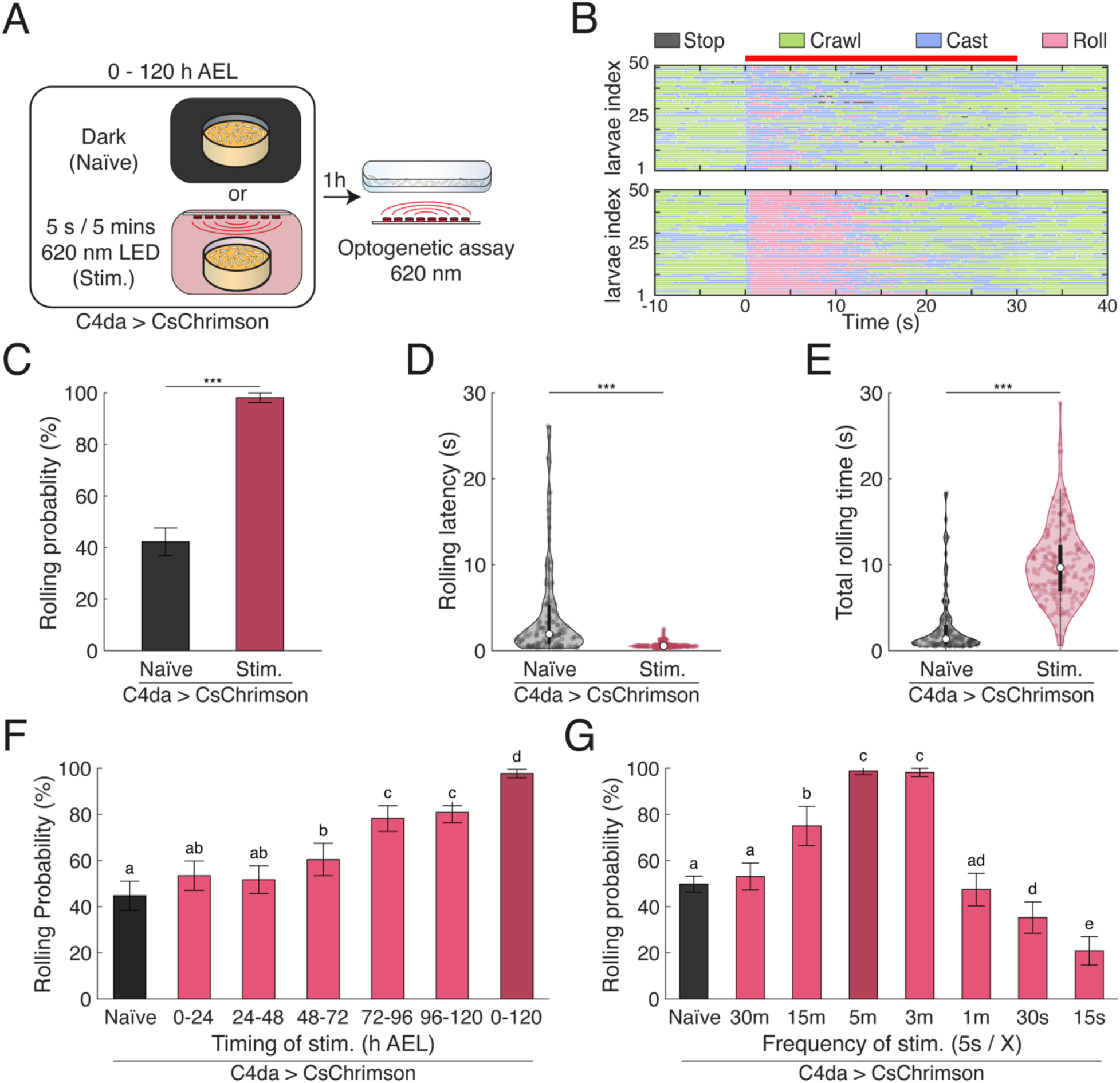
Noxious experience during development shapes nociceptive behaviours depending on timing and frequency. (A) Schematic of the experimental paradigm. Larvae were raised on food containing all-transretinal and allowed to develop for 120 hours until late third instar. Larvae were either kept in the dark (Naïve) or subjected to intermittent optogenetic activation of C4da nociceptive neurons during development (Stim.). Behavioural responses were tested via optogenetic stimulation at the late third instar stage. (B) Raster plots showing individual behavioural responses (stop, crawl, head cast, roll) over 30 s of optogenetic C4da stimulation. Each row represents a continuously tracked larva. Top - Naïve larvae. Bottom - Developmentally stimulated larvae. (C) Rolling probability over the 30 s stimulation window. Bars indicate mean; error bars denote 95% confidence intervals (n = 327, 207). Chi-square test, \*\*\**p* < 0.001. (D) Violin plot showing the rolling latency of larva during optogenetic stimulations (n = 138, 203). Welch’s ANOVA Test \*\*\**p* < 0.001. (E) Violin plot of total rolling time in response to optogenetic C4da stimulation (n = 138, 203). Welch’s ANOVA Test \*\*\**p* < 0.001. (F) Rolling probability in larvae based on developmental time window of stimulation. Bars show mean with 95% confidence intervals (n = 237, 234, 267, 187, 211, 283, 260). Chi-Square Test with compact letter display; groups not sharing a letter differ significantly at *p* <0.01. (G) Rolling probability as a function of stimulations during development. Error bars indicate the 95% confidence interval (n = 818, 277, 100, 174, 219, 194, 190, 168). Chi-square test with compact letter display; statistical significance at *p* < 0.01.

To investigate how the pattern of nociceptive sensory neuron stimulation during development influences nocifensive behaviour, we varied the stimulation parameters, including frequency, light intensity, and developmental timing. We first examined the effect of 24-hours stimulation across different developmental stages. Larvae stimulated during first instar stage showed a reduction in rolling latency compared to naïve control (Supp. Fig. 1A). Stimulation during the second instar stage further enhanced sensitization, leading to both decreased latency and increased rolling probability (Fig. 1F, Supp. Fig. 1A). Stimulation during the third instar resulted in the most pronounced sensitization, with significantly increased rolling probability, reduced latency and prolonged rolling duration compared to naïve animals (Fig. 1F, Supp. Fig. 1A-B). These findings indicate the behavioural sensitization can be robustly induced across developmental stages, though the extent of sensitization increases with larvae age. Next, we examined the impact of optogenetic stimulation intensity (applied between 96 - 120 h AEL) on nocifensive responses. Larvae exposed to higher LED intensities exhibited increased rolling probability, with behavioural effects reaching saturation at intensities above 0.35 µW/mm^2^ (Supp. Fig. 1C-E).

As noted above, previous studies have shown that continuous or high-frequency stimulation of nociceptive neurons can lead behavioural desensitization^35,43,44^. To further investigate this in our paradigm, we varied the frequency of C4da neurons activation while maintaining stimulation duration and intensity (5s pulses and 5.6 µW/mm^2^). As expected, high-frequency stimulation (every 15 or 30 seconds) significantly reduced rolling probability (20.8 ± 6.1% and 35.2 ± 6.8%, respectively), relative to naïve control (53.1 ± 5.9%). In contrast, lower-frequency stimulation (every 3, 5 or 15 minutes) induced significantly higher rolling probabilities (98.2 ± 1.8%, 98.9 ± 1.58%, 75.0 ± 8.5%, respectively) (Fig. 1G), with an apparent transition point around a stimulation interval of once per minute (47.4 ± 7.0%). Rolling latency and total rolling duration exhibited similar trends (Supp. Fig. 1F-G). These results demonstrate that behavioural outcomes are highly sensitive to the temporal pattern of nociceptive input. They suggest that distinct and potential opposing mechanisms underlies the encoding of sensitization versus desensitization evoked by prior experience.

### Behavioural sensitization reflects a decrease of the nociceptive threshold

The sensitization observed following optogenetic stimulation of C4da neurons during development could be an artifact of the technique rather than a biologically relevant process. If this was the case, optogenetically stimulated larvae might not show altered behavioural responses when presented natural noxious stimuli that also activate C4da neurons. To test this, we assessed the nocifensive response in both naïve and developmentally stimulated larvae following natural mechanical or chemical stimulation (Fig. 2A). When tested with mechanical stimulation, developmentally stimulated larvae were significantly more likely to exhibit rolling behaviour than naïve controls (66% vs. 38%; Fig. 2B). This increase in rolling was accompanied by reduction in the frequency of competing startle-like behaviour stopping (stimulated: 9% vs naïve: 25%) (Fig. 2B). Similarly, following exposure to noxious chemical stimulus (5% hydrochloric acid, HCl), optogenetically stimulated larvae rolled more frequently than naïve controls (93% vs. 79%, as calculated by proportion of animals rolling within 10 seconds; Fig. 2C). They also rolled with shorter latencies (3.27 ± 0.12 s vs 4.30 ± 0.15 s; Fig. 2C).

**Figure 2.**
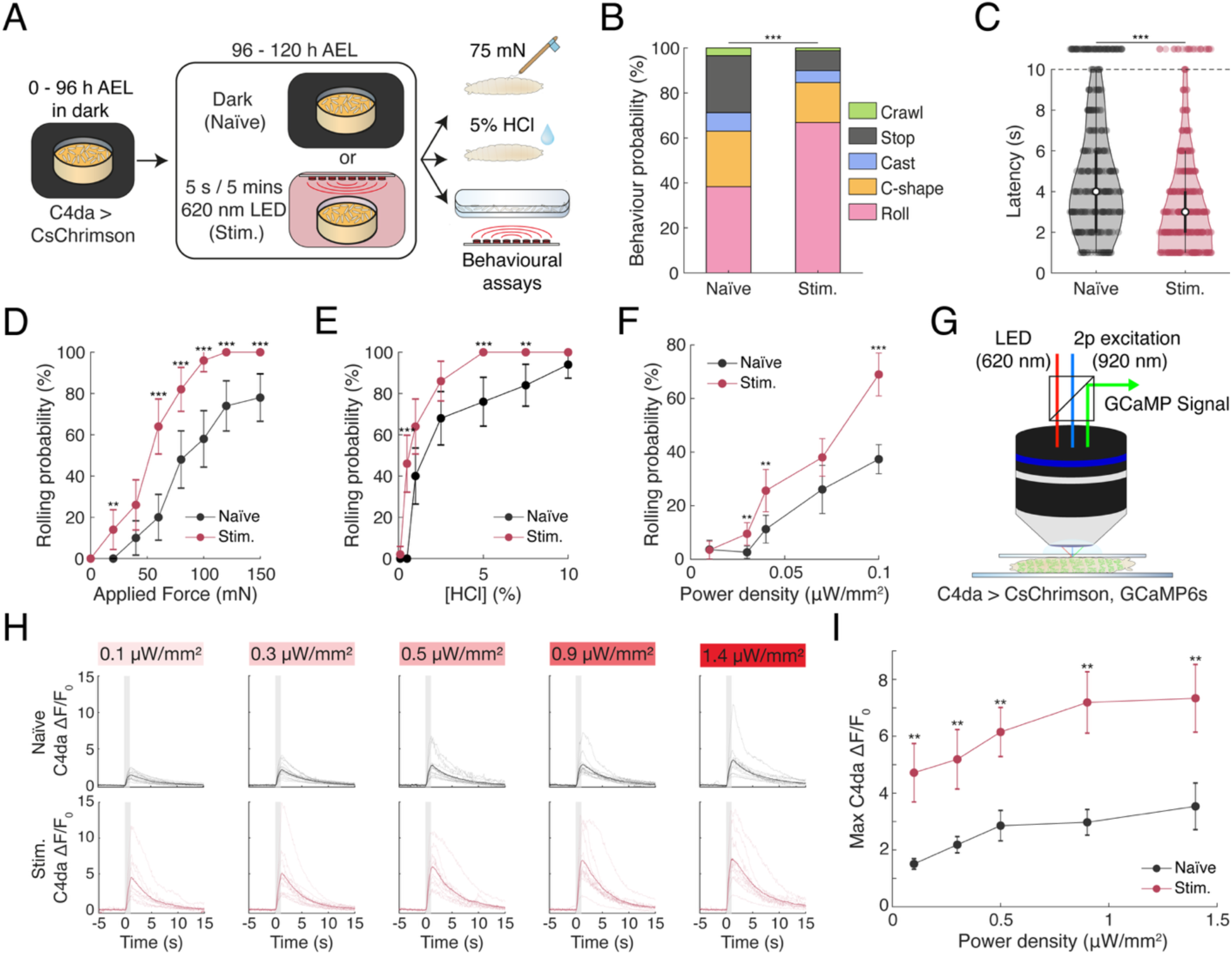
Prior noxious experience enhances larval nociceptive responses via increased C4da neurons excitability sensitivity. (A) Schematic of experimental design. Larvae were raised on all-trans retinal food, and developed in the dark until 96 hours post-collection. From 96h to 120h after collection, larvae were either kept in darkness (Naïve) or exposed to sporadic optogenetic activation of C4da neurons (Stim.). Nociceptive responses were then assessed using mechanical, chemical or optogenetic nociceptive assays. (B) Behavioural distribution in response to 75 mN mechanical stimulus, categorized into nociceptive responses (rolling and C-shape) and non-nociceptive (crawling, stopping, head-casting) responses (n = 146, 169). Mann-Withney U Test \*\*\**p* < 0.001. (C) Violin plot of latency to roll following application of 5% HCl (1.5µL). Larvae not responding within 10 s were plotted above the cutoff (n = 343, 350). Mann-Whitney U Test, \*\*\**p* < 0.001. (D) Dose-response curve for rolling probability as a function of force applied in the mechanical nociception assay. Error bars: 95% confidence interval (n = 50). Chi-square test, \*\**p* < 0.01, \*\*\**p* < 0.001. (E) Dose-response curve for the rolling probability as a function of the concentration in the chemical nociception assay. Error bars: 95% confidence interval (n = 50). Chi-square test, \*\**p* < 0.01, \*\*\**p* < 0.001. (F) Dose-response curve showing the rolling probability as a function of LED irradiance in optogenetic assays. Error bars: 95% confidence interval (n_Naïve_ = 110, 189, 142, 92, 305; n_Stim_ = 116, 189, 117, 184, 129). Chi-square test, ** *p* < 0.01, \*\*\**p* < 0.001. (G) Schematics of in vivo 2-photon imaging setup for mechanically restrained 3^rd^ instar larva (H-I) Calcium imaging of GCaMP6s signal in C4da soma during optogenetic C4da stimulation. n_Naïve_ = 12, 13, 12, 12, 11; n_Stim_ = 10, 12, 10, 11, 10. (H) Individual and average ΔF/F₀ response traces at varying irradiances. Shaded regions mark stimulation period; thick traces indicated group means. (I) Maximum ΔF/F_0_ across irradiances in naïve and stimulated animals. Error bars represent SEM. Mann-Whitney U Test ** *p* < 0.01.

These findings suggest that the increased rolling probability induced by optogenetic stimulation of C4da neurons may reflect a lowered threshold for nociceptive responses. If so, then dose-response curves for optogenetically stimulated versus naïve larvae should consistently show a leftward shift across different types of nociceptive stimuli. We first compared the mechanical thresholds between the two groups. At a typically subthreshold mechanical force (20 mN), 14 ± 9.6% of stimulated larvae exhibited rolling behaviour, whereas none of the naïve larvae responded. Stimulated larvae also showed significantly higher rolling probabilities at every points along the mechanical dose-response curve (40 mN:10 ± 8.3% vs 26 ± 12.2%; 60 mN: 20 ± 11.1% vs 64 ± 13.3%; 80 mN: 48 ± 13.8% vs 82 ± 10.6%; 100 mN: 58 ± 13.7% vs 96 ± 5.4%; 120 mN: 74 ± 12.2% vs 100% ; 150 mN: 78 ± 11.5% vs 100%) (Fig. 2D). We then tested chemical sensitivity using hydrochloric acid (HCl). Similar to mechanical stimulation, 46 ± 13.8% of the stimulated larvae rolled at concentrations that were subthreshold for naïve larvae (Fig. 2E). Additionally, stimulated larvae displayed higher probabilities across nearly all concentrations (1% HCl: 40 ± 13.6% vs 64 ± 13.3%; 2.5% HCl: 68 ± 12.9% vs 86 ± 9.6%; 5% HCl: 76 ± 11.8% vs 100%; 7.5% HCl: 84 ± 10.2% vs 100%), except for the highest tested concentration (10% HCl: 94% vs 98%) (Fig. 2E). Lastly, we assessed optogenetic stimulation. Developmentally stimulated larvae roll more frequently than naïve controls at the lowest light intensity (0.03 µW/mm^2^ – 2.65 ± 2.28% vs 9.52 ± 4.18%) and showed increased rolling probabilities at all higher intensities (0.04 µW/mm^2^: 11.3 ± 5.20% vs 25.6 ± 7.91%; 0.07 µW/mm^2^: 26.1 ± 8.97% vs 38.0 ± 7.01%; 0.1 µW/mm^2^: 37.4 ± 5.43% vs 69.0 ± 7.98%) (Fig. 2F).

Collectively, these data support the conclusion that developmental optogenetic stimulation of the C4da neurons induces behavioural sensitization by lowering the threshold for nociceptive responses across mechanical, chemical and optogenetic stimuli.

### Previous noxious experiences alter C4da neuron responses to stimuli, but not morphology

To understand the neural mechanisms underlying sensitization, we examined how developmental stimulation affect the activity of C4da sensory neurons. Larvae expressing both the calcium indicator GCaMP6s and optogenetic actuator CsChrimson in C4da neurons were subjected to acute optogenetic stimulation, and calcium responses were imaged in the dendritic field near the soma (Fig. 2G). Compared to naïve controls, developmentally stimulated larvae exhibited significantly higher calcium responses across all tested light intensities (*ΔF/F0*; naïve vs stim: 0.1 µW/mm^2^: 1.51 ± 0.19 vs 4.72 ± 1.03; 0.3 µW/mm^2^: 2.18 ± 0.29 vs 5.19 ± 1.05; 0.5 µW/mm^2^: 2.85 ± 0.53 vs 6.15 ± 0.86; 0.9 µW/mm^2^: 2.97 ± 0.45 vs 7.19 ± 1.08; 1.4 µW/mm^2^: 3.53 ± 0.82 vs 7.34 ± 1.19) (Fig. 2H-I and supplemental video 1 and 2). These results indicate that C4da neurons become more responsive following noxious stimulation during development.

Given the change in dendritic arborization are often associated with altered sensory neuron sensitivity^45,55–57^, we next examined whether morphological changes might account for the increased activity. We performed Sholl analysis on C4da dendritic arbors in developmentally stimulated (5 s / 5 min., 5.6 mW/mm^2^ for 96–120 AEL) and naïve larvae. This revealed that stimulation of C4da neurons during larval development did not drive changes in their arborization (Supp. Fig. 2). While the Sholl profile suggested a mild increase in branching in stimulated animals, these differences were not statistically significant (Supp. Fig. 2A–B). There were no significant differences in key parameters including the area under the Sholl curve (naïve vs stim.: 23.0 ± 1.24 vs 25.7 ± 1.32), critical radius (50.1 ± 3.4% vs 52.2 ± 4.1%), or number of intersections at critical radius (47.8 ± 2.8 vs 52.6 ± 3.6) (Supp. Fig. 2C-E). These findings indicate that developmental sensitization of C4da neurons is not mediated by structural change of their dendritic arbors. Instead, sensitization likely arises from functional changes driven by prior neuronal activity.

### The OAMB receptor in C4da neurons is necessary for experience-dependent sensitization

A previous study reporting behavioural desensitization following continuous noxious chemical stimulation of C4da neurons identified serotonin as a key neuromodulator mediating this effect ^43^. To investigate whether neuromodulators also play a role in the experience dependent sensitization observed in our experimental paradigm, we performed RNAi knockdown of various neuromodulator receptors specifically in C4da neurons. We then compared the behavioural responses of both naïve and developmentally stimulated larvae with and without receptor knockdown.

Our screen identified the octopamine receptor OAMB (Octopamine receptor in mushroom bodies) as a strong candidate involved in the sensitization of C4da neurons (Supp. Fig. 3A–C, Fig. 3A). RNAi knockdown of OAMB using two independent lines had minimal effect on nocifensive behaviour in naïve larvae, with only slight reduction or no reduction in rolling probability, rolling latency or total rolling duration compared to control naïve animals (Fig. 3A–C). However, in developmentally stimulated larvae, OAMB knockdown abolished the enhanced rolling response. Specifically, rolling probability was significantly reduced in OAMB knockdown animals compared to stimulated controls (No RNAi: 94.6 ± 2.64%; OAMB-RNAi-1: 36.0 ± 5.64%; OAMB-RNAi-2: 11.8 ± 3.11%) (Fig. 3A). In addition, latency to roll was increased (No RNAi: 0.87 ± 0.04 s; OAMB-RNAi-1: 2.85 ± 0.33 s; OAMB-RNAi-2: 5.50 ± 1.12 s), and the total rolling time was decreased (No RNAi: 6.76 ± 0.18 s; OAMB-RNAi-1: 3.05 ± 0.22 s; OAMB-RNAi-2: 1.43 ± 0.19 s) (Fig. 3B–C). These results demonstrate that OAMB is necessary in C4da neurons for experience-dependent sensitization prevents induced by developmental nociceptive stimulation. Knockdown of this octopamine receptor effectively blocks the increased behavioural sensitivity, suggesting that octopaminergic signaling acts directly on C4da neurons to mediate sensitization.

**Figure 3.**
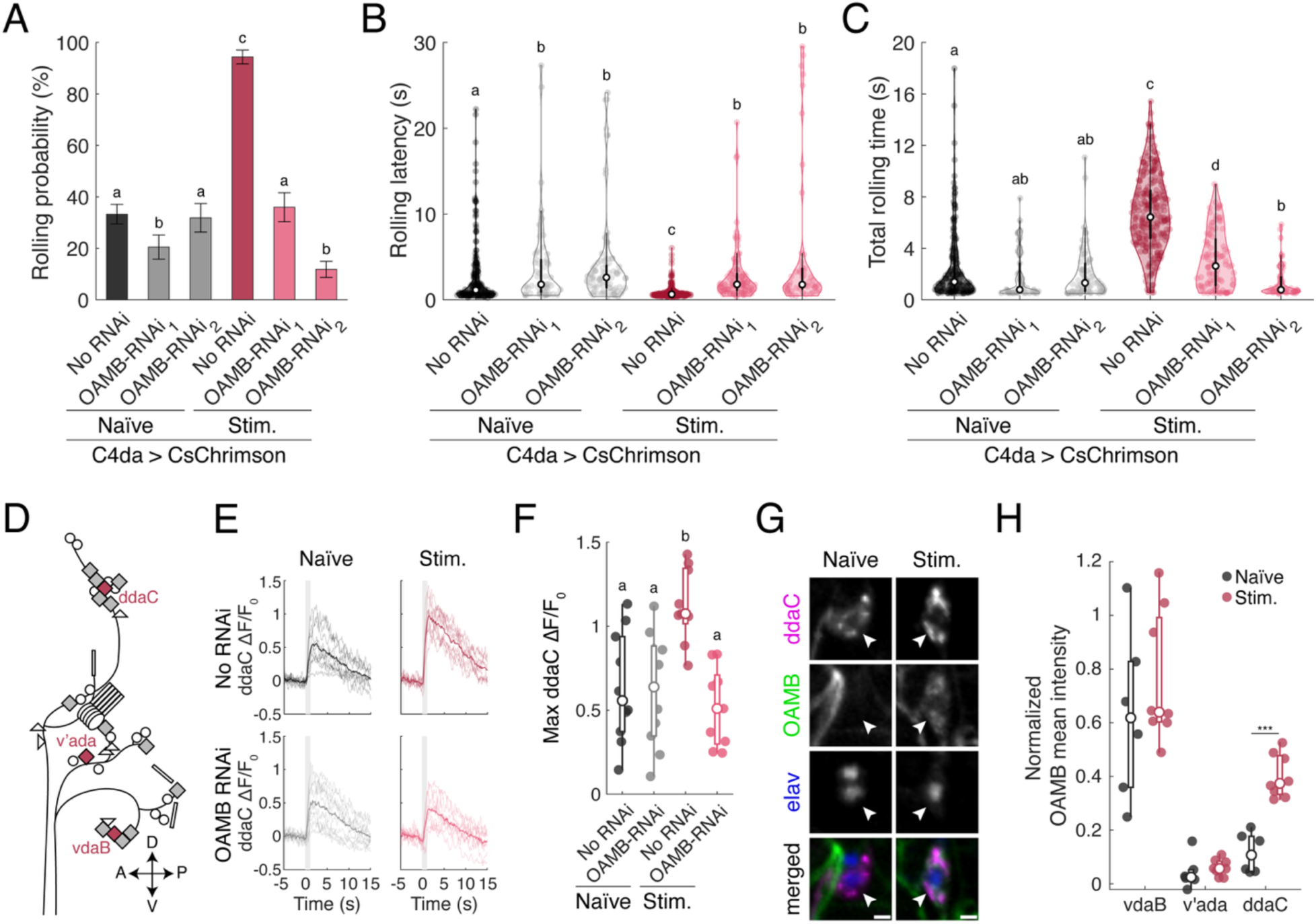
The octopamine receptor OAMB is required for experience-dependent sensitization to noxious stimuli. (A-C) RNAi knockdown of OAMB blocks the behavioural sensitization induced by prior noxious experience. (A) Rolling probability during optogenetic C4da stimulation. Error bars: 95% confidence interval (n = 589, 283, 270, 284, 278, 414). Chi-square test; compact letter display (CLD) shows statistical significance at *p* < 0.01. (B) Violin plot of rolling latency during optogenetic stimulation (n = 195, 58, 86, 268, 100, 49). Kruskall-Wallis followed by post-hoc Mann-Whitney test, CLD denotes *p* < 0.01. (C) Violin plot of total time each larvae spent rolling during optogenetic stimulations (n = 195, 58, 86, 268, 100, 49). Welch’s ANOVA with post-hoc Mann-Whitney test; CLD denotes *p* < 0.01. (D) Schematics of the larval somatosensory system, highlighting C4da neurons (red) (E-F) Calcium imaging of C4da neuronal responses during optogenetic stimulation with or without OAMB RNAi. n = 10, 10, 9, 9. (E) ΔF/F_0_ traces from individual neurons aligned to stimulus onset. Shaded region indicates stimulation period. (F) Maximum ΔF/F_0_ responses in naïve and stimulated larvae with or without RNAi. Mann-Whitney test; CLD indicates statistical significance at *p* < 0.01 (G-H) OAMB expression varies with neuronal identity and prior noxious experience. (G) Representative images mean OAMB expression in ddaC neurons in naïve and stimulated larvae. Scale bar = 10 µm. (H) Boxplot of normalized OAMB expression (relative to elav staining) across C4da neuron types and groups. (n_naïve_ = 6, n_stim_ = 8). One-way ANOVA Test \*\*\**p* < 0.001.

We next examined whether RNAi-mediated knockdown of OAMB affected the activity of C4da neurons *in vivo* by imaging calcium dynamics in the dorsal ddaC neuron using GCaMP (Fig. 3D). In naïve larvae, GCaMP signals in response to optogenetic activation of ddaC neurons were unaffected by OAMB knockdown (RNAi: 0.62 ± 0.11, no RNAi: 0.63 ± 0.10) (Fig. 3E and F). However, in developmentally stimulated larvae, GCaMP signals were significantly elevated in ddaC neurons with intact OAMB expression (1.12 ± 0.07), while this activity enhancement was abolished by OAMB knockdown (0.52 ± 0.08) (Fig. 3E and F). These results suggest that OAMB is required for the experience-dependent increase in sensory neuron activity following developmental stimulation.

To our knowledge, no prior studies have demonstrated OAMB expression in C4da neurons. To address this, we used a MiMIC-converted Trojan-GAL4 line targeting an intronic region shared by all OAMB isoforms^21^, to assess endogenous OAMB expression. We quantified co-expression of MiMIC-driven GFP with tdTomato-labeled C4da neurons proxy for OAMB expression in naïve and experienced larvae. GFP signal from the OAMB-trojan-GAL4 was consistently high in vdaB neurons, independent of experience (naïve: 0.75 ± 0.07, experience: .82 ± 0.06), but weak in ventral v’ada neurons (naïve: 0.30 ± 0.04, experienced: 0.40 ± 0.03) (Figure 3H, Sup Fig. 3D and E). In contrast, ddaC neurons exhibited negligible OAMB expression in naïve larvae (0.40 ± 0.03), but significantly elevated levels following developmental stimulation (0.63 ± 0.04) (Fig 3. G and H). These data indicate that while some C4da neurons (e.g. vdaB) express high levels of OAMB regardless of experience, ddaC neurons show experience-dependent upregulation of OAMB, suggesting a potential mechanism for observed sensitization.

### Octopaminergic signalling by tdc2+ neurons during noxious experience is necessary for nociceptive sensitization

Since OAMB in C4da neurons is required for experience-dependent sensitization, we surmised that the ligand of this receptor, octopamine, plays an essential role. To investigate how octopaminergic neurons contribute to the encoding of prior noxious experiences within the nociceptive system. We utilized the tdc2-GAL4 drive line to expression two RNAi lines targeting tyramine beta-hydroxylase (tbh-RNAi-1: JF02746, tbh-RNAi-2: HMS05829), a rate-limiting enzyme in octopamine synthesis. The tdc2-GAL4 driver recapitulates the expression pattern of the tyrosine decarboxylase 2 gene and has been shown to drive expression in both tyraminergic and octopaminergic neurons^58,59^. Consequently, this driver was previously used to study the role of octopaminergic neurons in larvae^58,59^.

Naïve larvae expressing tbh-RNAi-1 or tbh-RNAi-2 in C4da neurons displayed nocifensive behaviour comparable to those of naïve controls (Fig. 4A, Supp. Fig 4A-B). In contrast, stimulated larvae in which production of octopamine was knocked down by tbh-RNAi-1 or tbh-RNAi-2 showed reduced sensitization compared to stimulated controls, as assessed by the rolling probability (tbh-RNAi-1: 65.9 ± 5.02% vs control: 83.2 ± 3.91%; tbh-RNAi-2: 58.6 ± 5.72% vs control: 77.7 ± 5.61%), total rolling time (tbh-RNAi-1: 5.16 ± 0.31 s vs control: 8.52 ± 0.42 s; tbh-RNAi-2: 11.4 ± 0.56 s vs. control: 16.4 ± 0.55 s), and rolling latency (tbh-RNAi-1: 9.30 ± 0.35 s vs. control: 5.23 ± 0.26 s; tbh-RNAi-2: 3.29 ± 0.27 s vs control 0.84 ± 0.18 s) (Fig. 4A, Supp. Fig. 4A and B). These results support a model in which octopaminergic signalling from tdc2+ neurons contributes to the sensitization of C4da neurons.

**Figure 4.**
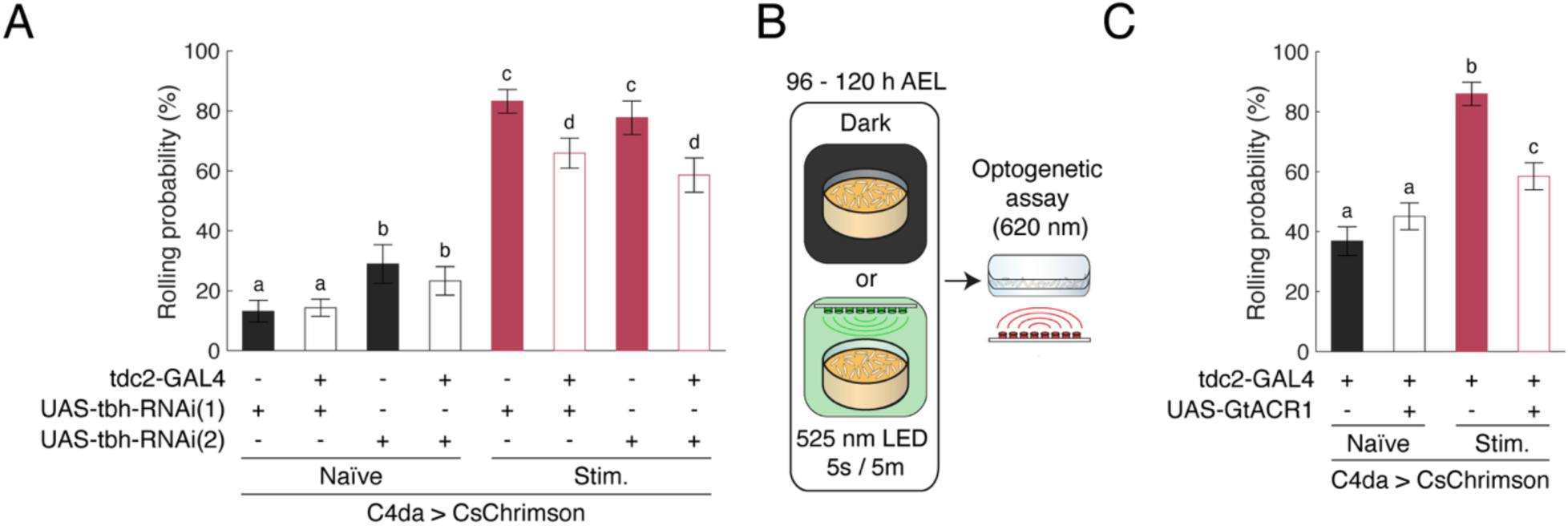
Octopaminergic signalling from tdc2+ neurons is required for experience-dependent nociceptive sensitization. (A) RNAi knockdown of tbh in tdc2+ neurons impairs experience-dependent sensitization without affecting baseline nociceptive responses. Rolling probability during optogenetic stimulation (n = 334, 579, 190, 300, 351, 343, 211, 285). Error bars represent 95% confidence interval. Chi-square test; CLD indicates statistical significance (*p* < 0.01). (B) Schematic of experimental design for inhibition of tdc2+ neurons during C4da neurons stimulation. 525 nm LED activate GtACR1 in tdc2+ neurons and CsChrimson in C4da neurons during development. Behavioural test was performed using 620 nm LED that only activate CsChrimson in C4da neurons. (C) Silencing tdc+2 neurons using GtACR1 alters nociceptive behaviour only in developmentally stimulated larvae. Rolling probability during optogenetic C4da neurons. (n= 391, 483, 306, 455). Error bars represent 95% confidence interval. Chi-square test; CLD indicates statistical significance (*p* < 0.01).

To further test the role of tdc2+ neurons in sensitization, we devised an experiment to transiently inhibit the activity of tdc2+ neurons during noxious stimulation. We expressed the green-light gated anion channel GtACR1 in tdc2+ neurons, while simultaneously expressing CsChrimson in C4da neurons. Larvae were then subjected to developmental stimulation using 525-nm (green) light (Fig. 4B). This wavelength effectively activates both GtACR1 and CsChrimson ^60,61^. Thus, we reasoned that a 525-nm light should supress tdc2+ neurons activity during optogenetic activation of C4da neurons. In contrast, stimulation with 620-nm (red) light should activate C4da neurons without affecting GtACR1 in tdc2+ neurons^60,61^. As expected, GtACR1 expression in tdc2+ neurons did not alter the probability of rolling in naïve larvae (control: 36.8 ± 4.8 %, GtACR1: 45.1 ± 4.4 %) (Fig. 4C). However, when tdc2+ neurons activity was inhibited during C4da neurons activation, we observed a significant reduction in sensitization in developmentally stimulated larvae (control: 86.0 ± 3.89 %, GtACR1: 58.5 ± 4.5 %) (Fig. 4C).

Since the activation of C4da during development results in both decreased response latency and increased total rolling time, we next assessed whether inhibition of tdc2+ neurons affect these parameters. Inhibiting tdc2+ neurons activity partially attenuated the reduction in rolling latency following stimulation (no GtACR1, naïve vs stim.: 4.24 ± 0.45 s vs 1.68 ± 0.15 s; with GtACR1, naïve vs stim.: 2.73 ± 0.36 s vs 1.25 ± 0.19 s – Cohen’s d = 0.760 vs 0.343). However, inhibition of tdc2+ neurons did not prevent the prolongation of the total rolling time (no GtACR1, naïve vs stim.: 3.40 ± 0.3 s vs 14.3 ± 0.5 s; with GtACR1, naïve vs stim.: 5.78 ± 0.34 s vs 18.6 ± 0.50 s), (Supp. Fig. 4C-D). Together, these finding demonstrate that octopaminergic signalling during noxious stimulation is critical for experience-dependent sensitization, consistent with the results observed using tbh-RNAi. Specifically, activity of tdc2+ neurons during harmful experience is required for the increase in rolling probability and the reduction in response latency, but not for the prolongation of the overall nocifensive behaviour.

### Octopamine is sufficient to induce sensitization of the nociceptive system

Having shown that octopaminergic activity is necessary for sensitization, we next investigated whether octopamine itself is sufficient to induce sensitization in larval nociception. To test whether octopamine could mimic effect of nociceptive stimulation during development, we supplemented larvae food with octopamine at concentrations previously shown to rescue phenotypes in octopamine-deficient larvae^18,62^. In line with prior studies^18,62^, and to minimize potential acclimation to the additive, larvae were fed octopamine-enriched food for two hours prior to nociceptive optogenetic, mechanical, or chemical assay (Fig. 5A). Using optogenetic stimulation, we found that dietary octopamine significantly increased the rolling probability at all tested concentrations (5 mM: 69.1 ± 6.15%, 25 mM: 72.3 ± 6.07%, 50 mM: 77.0 ± 6.25%) compared to untreated control (0 mM: 49.0 ± 7.03%) (Fig. 5B). Since 5mM was sufficient to elicit behavioural changes, we used this octopamine concentration for subsequent experiments. In the mechanical stimulation assay, larvae exposed octopamine exhibited increased the rolling probability (52%) compared to untreated control (31%) (Fig. 5C). Similarly, in the chemical stimulation assay, octopamine-fed larvae showed an elevated (98%) rolling probability relative to control (92%) and reduced rolling latency (control: 3.16 ± 0.18 s; 5mM 2.53 ± 0.14 s) (Fig. 5D).

**Figure 5.**
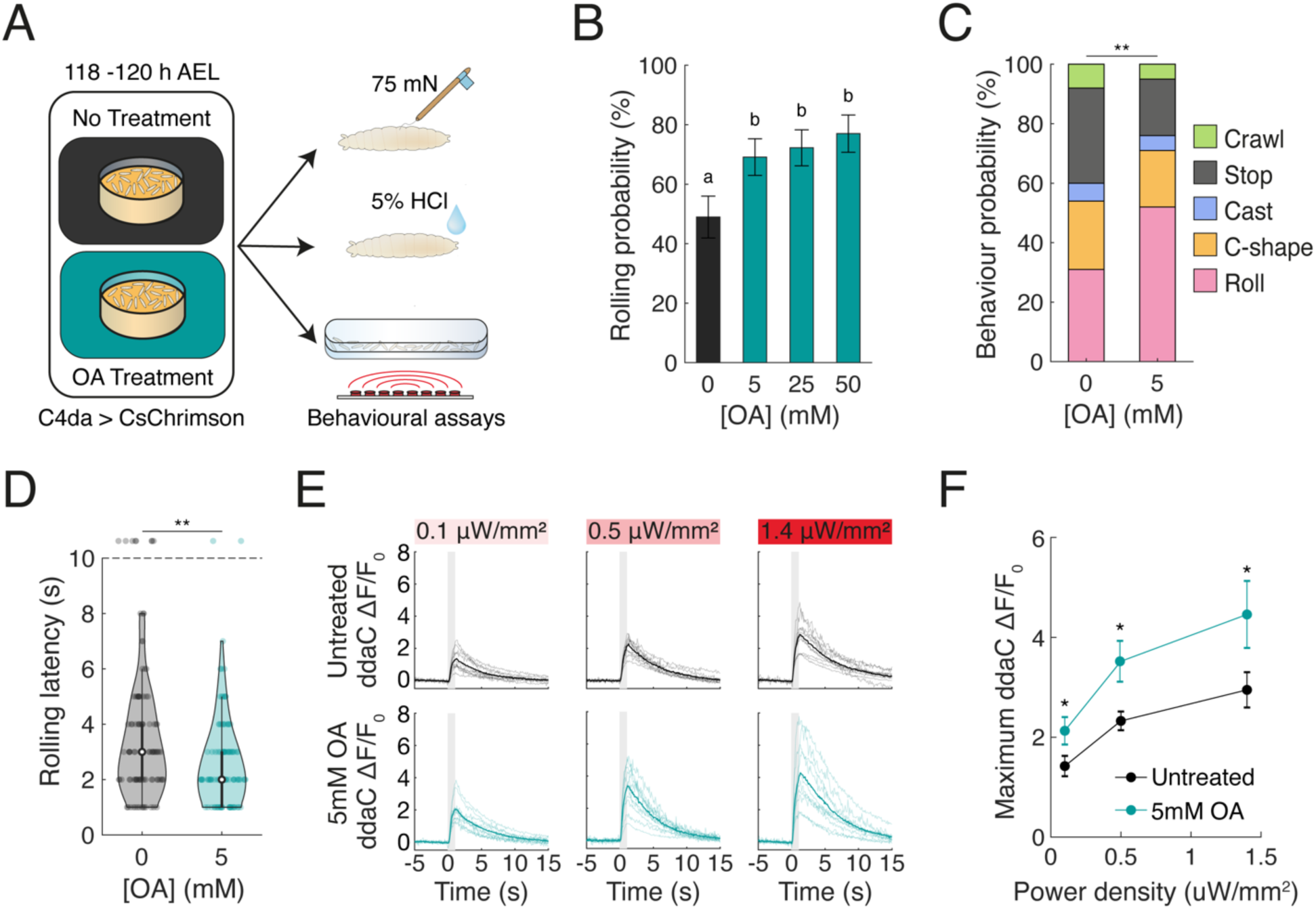
Octopamine is sufficient to enhance nociceptive response. (A) Schematics of experiment. Larvae were fed food supplemented with octopamine at varying concentrations prior to behavioural assays. (B-D) Octopamine feeding increases rolling in response to real and fictive nociceptive stimuli. (B) Rolling probability over the 30 s optogenetic stimulation of C4da neurons. Bars indicate mean; error bars denote 95% confidence intervals (n= 194, 217, 209, 174). Chi-square test; CLD indicates statistical significance (*p* < 0.01). (C) Behavioural distribution in response to 75 mN mechanical stimulus (n= 100/group). Mann-Whitney U Test, \*\**p* < 0.01. (D) Rolling latency following 5% HCl exposure (n= 100/group). Mann-Whitney U Test, \*\**p* < 0.01. (E-F) Octopamine feeding enhances C4da neuronal responses to optogenetic C4da stimulation. (E) Calcium traces from individual C4da neurons. Shaded regions denote stimulus period: bold line show group means. (n_Untreated_ = 10, 10, 10; n_Treated_ = 10, 12, 10) (F) Maximum ΔF/F0 in untreated and treated larvae. Mann-Whitney U Test \**p* < 0.05.

These data support the idea that octopamine enhances larval responses to noxious stimuli. This effect could result from changes in sensory neurons sensitivity, or from alterations in motor neurons where octopamine is known to regulates locomotion by modulating motor neuron activity through a balance between tyramine and octopamine^18,23,63^.To determine whether the behavioural effects of octopamine were at least partially derived from changes in sensory processing, we examined the calcium responses in C4da neurons in whole-mount larvae following octopamine treatment. Consistent with responses observed in larvae previously exposed to noxious stimulation, octopamine-fed larvae exhibited significantly enhanced calcium activity in C4da neurons upon optogenetic stimulation (Fig. 5E-F). Collectively, these results suggest that the octopamine sensitizes C4da neurons that likely underlies the observed behavioural changes.

### Ventral unpaired median neurons (VUMs) are critical for experience dependent sensitization

In *Drosophila* larvae, approximately ∼100 octopaminergic neurons are present in central nervous system (CNS)^64,65^. Octopaminergic neurons found in the ventral nerve chord (VNC) can be categorized in distinct classes of neurons, including the ventral unpaired median neurons (VUMs), a class of type II motor neuron, the dorsal unpaired median neurons (DUMs), the abdominal leucokinin producing neurons (ABLKs) and the ventral paired median neurons (VPMs)^64–66^. To investigate anatomical relationship between octopaminergic and C4da sensory neurons, we first employed immunohistochemistry to compare the morphology of tdc2+ neurons and C4da neurons. This analysis revealed that VUM neurons extend processes that overlap with the both anterior and posterior regions of C4da neurons axon terminal (Fig. 6A-B).

**Figure 6.**
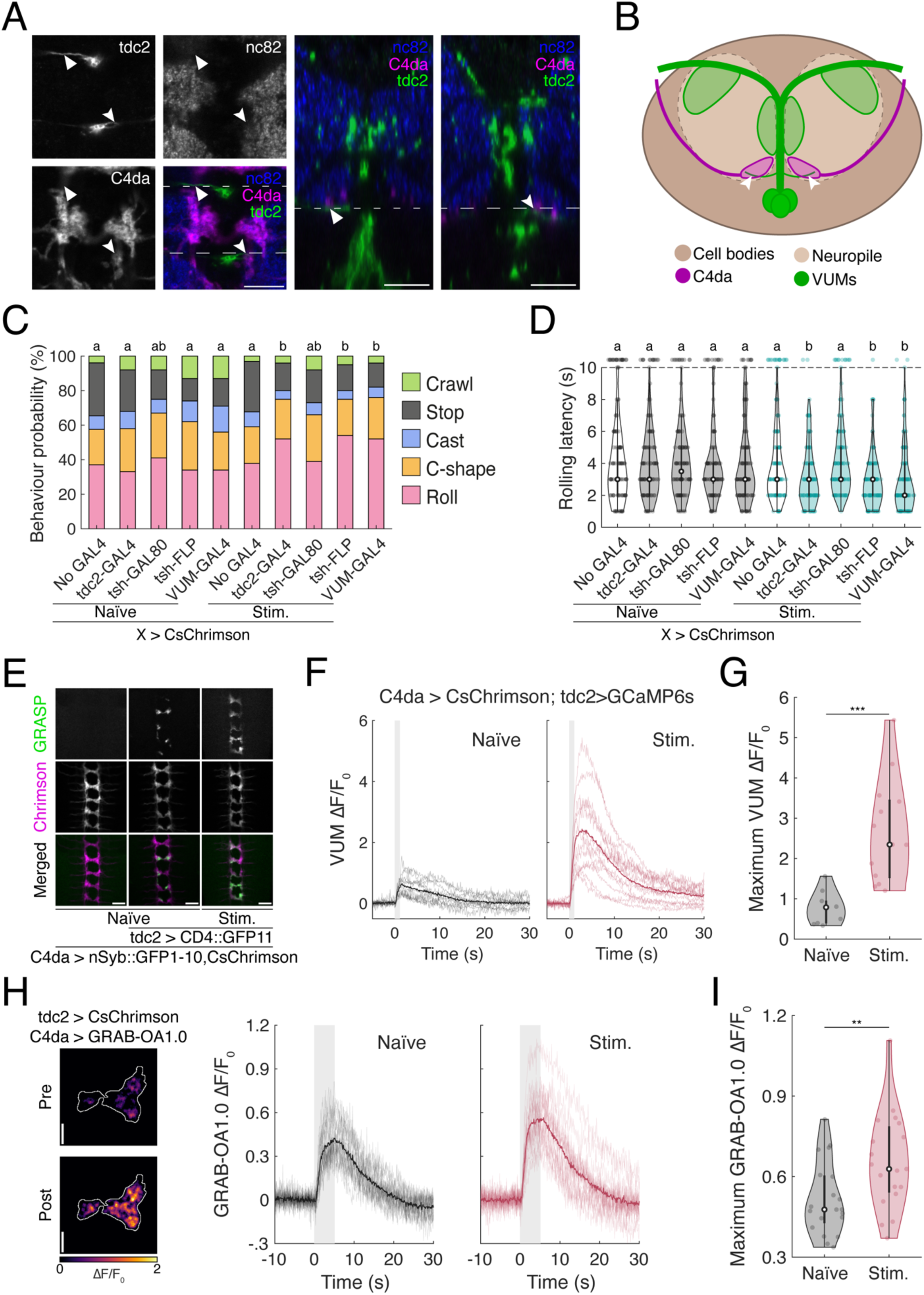
Octopaminergic VUM neurons mediate experience-dependent sensitization in the nociceptive circuit. (A) Expression pattern of VUM neurons and C4da neurons. Immunostaining of VUM neurons and C4da neurons. Arrow heads and triangle indicate the intersection between the abdominal VUM arbor and the C4da axon terminal. Scale bar = 20 µm. (B) Diagram summarizing the anatomical relationship between VUM neurons and C4da neurons. (C-D) Stimulation of VNC-restricted tdc2⁺ neurons recapitulate the sensitization induced by full tdc2⁺ neuron activation. (C) Behavioural distribution in response to 75 mN mechanical stimulus (n = 205, 100, 100, 100, 100, 198, 100, 100, 100, 100). Kruskall-Wallis test followed by post-hoc pairwise Mann-Whitney test; CLD indicates statistical significance at *p* < 0.01. (D) Violin plot of latency to roll following application of 5% HCl (n = 113, 100, 100, 100, 100, 112, 100, 100, 100, 100). Kruskall-Wallis test followed by post-hoc pairwise Mann-Whitney test; CLD indicates statistical significance at *p* < 0.01 (E) Developmental stimulation strengthens the synaptic connection between C4da neurons and tdc2⁺ neurons. Representative image of trans-synaptic GRASP signal between C4da and tdc2⁺ neurons. Scale bar = 20 µm. (F-G) Calcium imaging of tdc2⁺ VUM neurons reveals enhanced activation by C4da input after noxious developmental stimulation. (F) GCaMP6s traces of individual and average response in VUM neurons; shaded region indicates stimulus. (G) Peak ΔF/F₀ in naïve vs. stimulated larvae (n= 10, 13). Mann-Whitney test *** *p* < 0.001. (H-I) GRAB-OA1 imaging of C4da neurons in response to tdc2+ neurons simulation. (H) Heatmap represent average intensity of GRAB-OA signals in C4da axon terminal before and after stimulation in a naïve animal (left panel). GRAB-OA traces of individual and average C4da neurons; shaded region indicates stimulus. (H) Maximum ΔF/F_0_ in naïve and stimulated larvae (n= 20, 19). One-way ANOVA, ** *p < 0.01*.

To further test the role of VUMs in experience-dependent sensitization, we used targeted genetic approaches to manipulate CsChrimson expression. Specifically, we used tsh-LexA to drive FLP or GAL80, enabling us to restrict CsChrimson expression to either brain or the VNC (Supp. Fig. 5A-C). In parallel, we used the VUM-GAL4 line (GMR46B08-GAL4), which selectively targets expression of CsChrimson to VUM neurons and small subset of secondary abdominal neurons (Supp. Fig. 5D). Larvae expressing CsChrimson in the targeted neuronal populations for 24 hours (5 seconds every 5 minutes), and behavioural responses were compared the results to control groups: larvae lacking GAL4 (no CsChrimson expression) and larvae expressing CsChrimson in all tdc2+ neurons. We hypothesized that if the tdc2+ neurons activity is critical for nociceptive sensitization, their stimulation would mimic the effect of C4da neurons stimulation. Moreover, if VUM neurons are the primary mediators of this effect, then stimulating them in isolation should be sufficient drive similar behavioural sensitization. Supporting this hypothesis, optogenetic activation of tdc2+ neurons in VNC neurons alone significantly increased the rolling probability in response to a mechanical stimulus, relative to naïve controls or larvae lacking CsChrimson expression (Fig. 6C). A similar effect was observed in response to noxious chemicals, with treated larvae displaying both reduced rolling latency and increased rolling probability (Fig. 6D). These effects were comparable to those observed in larvae when all tdc2+ neurons were activated (Fig. 6C-D). In contrast, activation of tdc2+ neurons in the brain did not significantly alter nociceptive responses (Fig. 6C-D). Taken together, these findings strongly suggest that activation of VUM neurons activation is sufficient to drive experience-dependent sensitization of nociceptive behaviour. This likely occurs through octopaminergic modulation of C4da neurons.

Having established that VUMs activity is critical for nociceptive sensitization, we next investigated the functional relationship between VUMs and C4da neurons, both in naïve and experienced larvae. To assess potential anatomical connectivity, we employed activity-dependent GFP reconstitution across synaptic partner (syb-GRASP) ^67^, assuming that C4da neurons function as presynaptic partner to VUMs. In line with our anatomical descriptions (Fig. 6A-B), naïve animals displayed sparse but consistent GRASP signals, most reliably in segments T2, A2-A4 and A8/9 (Fig. 6E, Supp. Table 1). To further evaluate functional connectivity, we imaged calcium responses in VUM neurons following optogenetic activation of C4da neurons. While naïve larvae exhibited moderate calcium transients (average max ΔF/F0: 0.76 ± 0.13), experienced larvae displayed significantly larger responses (2.67 ± 0.36), indicating that prior noxious stimulation enhances synaptic transmission from C4da neurons to VUMs (Fig. 6F-G).

To explore whether this interaction is unidirectional or bidirectional, we activated tdc2+ neurons using CsChrimson and monitored octopaminergic transmission in C4da neurons using octopamine sensor GRAB-OA1.0^15^. This analysis revealed that C4da axon terminals receive octopaminergic input upon activation of tdc2+ neurons, and this input is significantly elevated in larvae with prior activation of the tdc2+ neurons (Fig. 6H-I). Together, these findings support the existence of positive feedback loop between C4da sensory neurons and VUMs octopaminergic neurons. In this circuit, noxious stimulation strengthens synaptic connectivity and mutual responsiveness: C4da neurons more effectively activate VUMs, while VUMs, in turn release greater amounts of octopamine onto C4da neurons in experienced larva. This bidirectional enhancement provide to amplify nociceptive sensitization following developmental experience.

## Discussion

In this study, we demonstrate that developmental activation of nociceptive C4da neurons in *Drosophila* larvae leads to sensitization of nociceptive behaviour. We show that this experience-dependent sensitization is accompanied by enhanced activity of C4da neurons in response to acute stimulation but occurs without significant changes in dendritic morphology. We identify the octopamine receptor OAMB as a key molecular mediator of this sensitization, showing that its expression in C4da neurons is required for the enhancement of behavioural and neural responses. Furthermore, we reveal that OAMB is expressed in C4da neurons in an experience-dependent manner, particularly in ddaC neurons, indicating dynamic regulation of receptor expression. We further identify VUM octopaminergic neurons as both anatomically and functionally connected to C4da neurons, and show that activation of VUMs is sufficient to induce sensitization. Finally, we provide evidence for a positive feedback loop between C4da neurons and VUMs, as prior nociceptive experience strengthens the synaptic and functional connectivity between these populations, leading to increased responsiveness of both partners and amplification of nociceptive sensitization.

Our findings highlight a form of neuromodulator-driven plasticity within the *Drosophila* larvae nociceptive circuit in which prior noxious experience alters both the sensory neurons and their associated modulatory inputs. Specifically, we show that stimulation of C4da nociceptive neurons during development leads to an upregulation of the octopamine receptor OAMB, enhanced octopaminergic signaling, and increased responsiveness in both C4da neurons and their upstream VUM modulatory neurons. These results suggest that the circuit adaptation involves changes at multiple nodes–both pre- and postsynaptic changes–enabling a more robust behavioural response to future noxious stimuli. This plasticity is experience-dependent and tightly linked to the internal state of the animal, supporting the idea that neuromodulatory systems can encode and store information about past sensory experiences to shape future behaviour^7,10–13,20,22,23^. Such mechanisms are reminiscent of central sensitization phenomena in vertebrates, where prior injury or repeated noxious input leads to persistent hypersensitivity and exaggerated behavioural responses ^50,68^. Rather than being confined to a single locus, this form of plasticity appears to rely on reciprocal adjustments across sensory and modulatory components of the circuit. These bidirectional changes ensure that information about past noxious experiences is integrated into both receptor expression and circuit connectivity, supporting flexible yet precise tuning of nociceptive sensitivity that balances protective responsiveness with the need to avoid maladaptive overreaction.

A growing body of evidence points toward a neuromodulatory and immune mechanisms by which *Drosophila* larvae dynamically regulate nociceptive behaviour based on the temporal pattern that is used to activate different classes nociceptive neurons ^35,43,44^. Previous work has shown that serotonin contributes to behavioural desensitization following persistent, high-frequency stimulation of C4da nociceptive neurons, suggesting a dampening mechanism engaged by sustained noxious exposure^43^. In contrast, recent results demonstrate that octopamine mediates sensitization, enhancing nociceptive responsiveness following intermittent, low-frequency developmental stimulation. Preliminarily, our data suggest that octopaminergic neurons are preferentially responsive to low-frequency input, while serotonergic neurons may be tuned to continuous, high-frequency stimulation. This dichotomy implies a frequency-dependent switch in neuromodulatory state that governs opposing behavioural outcomes—sensitization or desensitization—depending on the nature of the threat. Such a mechanism could enable the larval nervous system to implement context-specific adaptations: brief, sporadic threats may trigger heightened vigilance via octopaminergic sensitization, while prolonged or inescapable stimuli may engage serotonergic desensitization pathways to prevent maladaptive overreaction. This model reflects analogous processes in vertebrates, where distinct neuromodulatory systems mediate either pain amplification or suppression depending on stress duration and intensity^69^. By integrating frequency-dependent sensory coding with neuromodulatory feedback, *Drosophila* may employ a flexible system that adjusts behavioural responses to both immediate threat and cumulative sensory history—supporting a form of state-dependent plasticity in nociceptive processing (see *Upreti et al., 2019*^70^, for a similar dual-process mechanism in *Aplysia*, involving structural plasticity in neuromodulatory circuits alongside synaptic changes between sensory and downstream neurons).

The identification of an octopaminergic feedback loop modulating nociceptive sensitivity in *Drosophila* larvae raises compelling questions about the evolutionary conservation of neuromodulatory control over pain and arousal states. Octopamine, the invertebrate analog of norepinephrine, serves many overlapping functions with its vertebrate counterpart—modulating arousal, locomotion, aggression, learning, and now, as we show, nociceptive plasticity. Our finding that the *Drosophila* OAMB receptor, a putative homolog of the vertebrate α1-adrenergic receptor^71^, is both expressed in nociceptive sensory neurons and necessary for sensitization suggests that direct neuromodulatory input onto primary nociceptors may be a deeply conserved feature of bilaterian nervous systems. In mammals, peripheral adrenergic signaling has been implicated in pain hypersensitivity, with α1-adrenoceptors shown to enhance nociceptor excitability and contribute to chronic pain states^72–75^. Similarly, the feedback loop we propose in *Drosophila*—where prior nociceptive experience amplifies neuromodulator release and receptor expression in nociceptors—parallels mechanisms observed in vertebrates, where injury or stress can lead to increased adrenergic tone and receptor upregulation in the periphery^72–77^. These similarities hint at a shared evolutionary strategy in which stress-related neuromodulators function as gain controls for sensory systems, tuning behavioural output to match the organism’s internal state and environmental context. Our findings in *Drosophila* not only illuminate fundamental principles of nociceptive plasticity in invertebrates, but also provide a genetically tractable model to study how neuromodulatory systems interface with sensory circuits to shape adaptive—and potentially maladaptive—pain-related behaviours across species.

## Methods

### Fly stocks and maintenance

The fly stocks carrying the following genetic reagents were used in this study. The following stocks were obtained from the Bloomington Drosophila Stock Center (BDSC): UAS-GRAB(OA1.0) in attP40 (BDSC #604600); LexAop2-CsChrimson.tdTomato in VK00005 (BDSC #82183); UAS-IVS-CsChrimson.mVenus in attP18 (BDSC #55134); UAS-IVS-CsChrimson.mVenus in attP2 (BDSC #55136); LexAop-GAL80 in attp40 (BDSC #32214); GMR27H06-LexA in JK22C (BDSC #94664); GMR46B08-GAL4 in attP2 (BDSC #47361); LexAop-nSyb-spGFP1-10, UAS-CD4-spGFP11 (BDSC #64315); ppk-CD4::tdTomato (BDSC #35844); tdc2-GAL4.S in attP2 (BDSC #52243); tdc2-lexA::p65 in attP40 (BDSC #52242); TRiP.HMS05829 in attP40 (BDSC #67968); TRIP.JF01673 in attP2 (BDSC #31171); TRIP.JF01732 in attP2 (BDSC #31233); TRIP.JF02746 in attP2 (BDSC #27667); UAS-Dcr2.D in 2 (BDSC #24650); UAS-GtACR1.d.EYFP in attP2 (BDSC #92983). The following stocks were kindly gifted by G. Rubin: UAS-IVS-mCD8::RFP in attP18; UAS-IVS-myr::GFP in attp2; UAS-Syn21-Chrimson88-tdTomato-3.1 in attP18; LexAop-CsChrimson-tdTomato in attP18; LexAop-mCD8::GFP in su(Hw)attP8; UAS-dsFRT-Chrimson in attp18; UAS-Syn21-opGCaMP6s in su(Hw)attP8; LexAop-FLP in attp40. Tsh-LexA and ppk1.9-LexA were kind gifts from J. Simpson. ppk1.9-GAL4 was kindly gifted by D. Tracey. UAS-IVS-GCaMP6s in attp2 was a kind gift from J. Jayaraman. Mi{Trojan-GAL4.1-Oamb}[MI12417-TG4.1] was a kind gift from R. S. Stowers. Information about the genetic constructs carried for experimental purposes can be found in supplemental table 2.

All *D. melanogaster* lines used in this study were maintained on standard cornmeal medium (Bloomington Drosophila Stock Center formulation) in a temperature- and humidity-controlled incubator at 25 °C under a 12h light-dark cycle. For optogenetic experiments, flies and larvae were reared on same medium supplemented with 0.2 mM of all-trans retinal (Toronto Research Chemicals, R240000), prepared fresh and protected from light.

#### Developmental optogenetic stimulations

Embryos were collected every 24 hours on food plates supplemented with all-trans retinal and larvae were reared in the dark until the third instar stage in the dark. To activate neurons during development, larvae expressing CsChrimson were exposed to pulsed red light stimulation (wavelength - 620 nm). Larvae were housed under LED array programmed to deliver five seconds pulses every five minutes at a power density of 3.53 µW/mm^2^ unless otherwise noted. Light delivery was controlled via Raspberry Pi running on Raspian OS, using a Python script controlled the array with the pigpio library general-purpose input-output control.

#### Octopamine treatment

To manipulate octopaminergic signaling, larvae were fed octopamine hydrochloride (≥95% purity, Sigma Aldrich), following protocols adapted from *Saraswati et al., 2004* ^18^ and *Chen et al., 2013* ^62^. Briefly, molten cornmeal agar was supplemented with red food coloring and octopamine at indicated concentrations. Two hours prior to the behavioural testing, larvae were transferred to octopamine-containing plates or dye-only control plates. Only larvae that exhibited visible red coloration in the abdomen―indicating ingestion of the octopamine-enriched food―were selected for testing.

### Behavioural assays

Prior to behavioural testing, larvae were separated from the food medium using a 15% sucrose solution, transferred to a mesh sieve, and gently rinsed with distilled water. Larvae were then transferred to the appropriate behavioural arena, as described below. All behavioural manual annotation were conducted blind to genotype to minimize bias.

#### Mechanical nociception

Mechanical nociception assays were conducted as described in *Zhu, Boivin et al., 2023* ^78^, as adapted from *Hu et al., 2017* ^33^. Briefly, groups of 5-7 third-instar larvae were placed on humidified petri dish. Two subsequent mechanical stimuli were delivered to the dorsal side of each larva (segments A4-A6) using calibrated custom-made Von Frey filament (20-150mN). Behavioural responses were scored according to predefined ethogram on scale from 1 to 5, with 1 indicating no observable response and 5 indicating full rolling.

#### Chemical nociception

Chemical nociception was assessed using methods adapted from *Lopez-Bellido et al., 2019*^25^. Group of 5-7 larvae were placed on a petri dish and briefly dried with a Kimwipe. Individual larvae were then exposed to 1.5 μl of hydrochloric acid (HCl; 0.5%-10% concentration) applied to the tail using a pipette. Response latency was recorded, with the first complete rolling behaviour considered the response. Larvae failing to roll within 10 s were classified as non-responders. Thermal nociception

Thermal nociception assays followed the protocol by *Zhong et al., 2010* ^79^. Group of 5-7 larvae were placed on a humidified petri dish. A calibrated heat prob was applied to the dorsal side between segments A4 and A6. Response latency was measured as the time to the first complete rolling behaviour. Larvae that did not roll within 10 s were considered non-responders.

#### Optogenetic apparatus and assays

Optogenetic assays were adapted from *Ohyama et al., 2013* ^80^, with modifications to the hardware configuration as described by Zhu et al., 2024 ^40^. The experimental setup consisted of a C-MOS Camera (BlackflyUSB3, BFLY-U3-23S6M-C, FLIR) with fixed focal length lens (Edmund, 56-529), 750 nm long-pass filter (Edmund, 66-575) and 850-nm infrared LED illumination for tracking, a 620-nm LED array for optogenetic stimulation, and a dedicated computer for stimulus delivery and data acquisition. Larval behaviour was recording and analysed using the Multi Worm Tracker (MWT), with stimulus delivery controlled via an integrated stimulus module.

For each trial, larvae were placed evenly on a 25cm x 25cm square agar plate (2% agar, pre-humidified using 15 ml of de-ionized water). The stimulation protocol consisted of two cycles of 30 s red-light stimulation (100 Hz, 90 % duty cycle) interleaved with 30 s rest periods. Unless otherwise noted, experiments were conducted at a power density of 0.84 µW/mm^2^.

### Behavioural data analysis

#### Behaviour detection

Behavioural feature extraction from optogenetic experiments was performed using the tracking data collected by the Multi-Worm Tracker (MWT) software. This data included larval contour, spine, and center of mass coordinates over time for individual animals. Kinematic parameters—including larval length, width, body area, speed, crabspeed (lateral movement against body axis), body curvature, head angle, and directional bias—were computed using the Choreography software suite, as described in *Ohyama et al., 2013*^80^, and in *Ohyama et al., 2015*,^30^. Larvae were included in the analysis only if they were continuously tracked for a minimum of 5 seconds and moved at least one body length. Collisions between larvae led to termination of tracking for the involved individuals, after which new object IDs were assigned.

Behavioural classification was made using machine-learning approaches applied to the features computed by the Choreography software. Non-nociceptive behaviours (i.e., crawling, hunching, stopping, and non-descript head casting) relied on a classifier developed and described by *Masson et al., 2020*^54^. Nociceptive behaviour, specifically rolling behaviour was detected using a classifier developed through the Janelia Automatic Animal Behaviour Annotator (JAABA)^30,53^, trained using the rolling behaviour described in *Zhu et al., 2024*^40^. To improve the specificity of rolling detection, events lasting ≥ ≥0.5 s are considered as valid rolling behaviour.

#### Behaviour quantification

Detected behavioural events were aggregated and quantified using MATLAB scripts. Only larvae that were successfully tracked throughout the entire stimulation period were included in the analysis. For rolling behaviour, the following metrics were computed:

Rolling probability: the proportion of larvae that exhibited at least one rolling event during the stimulation period, relative to the total number of animals recorded.

Rolling latency: the time elapsed from the onset of the stimulus to the initiation of the first rolling event.

Total rolling time: the cumulative duration of all rolling events within the stimulation window.

Larvae that did not exhibit rolling behaviour during the stimulation were excluded from rolling latency and total duration analysis to avoid skewing comparisons due to differing response rate across groups.

### Larval dissections and immunohistochemistry

Dissections and immunohistochemistry were conducted using standard protocols as described by *Patel, 1994*^81^. Briefly, either larval CNS or larvae fillet with intact peripheral nerve system (PNS) and CNS were dissected in phosphate-buffered saline (PBS). Tissues were then fixed in 4% paraformaldehyde (PFA) in PBS for 20 minutes at room temperature (RT), followed by three washes in PBS three times and two washes in PBS containing 0.4% Triton-X (PBX). Samples were then blocked in PBX supplemented with 5% normal goat serum (NGS) for 1 hour at RT. Following blocking, samples were incubated in primary antibodies for 1 hour at RT and subsequently overnight at 4°C. The following primary antibodies were used in PBX: chicken anti-GFP (1:3000, ab13970, Abcam), mouse anti-GFP (1:500, G6539, Sigma), rabbit anti-dsRed (1:1000, 632496 Takara/Clontech), rabbit anti-tdc2 (1:2000, ab128225, Abcam), mouse anti-nc82 (1:50, DHSB), rat anti-elav (1:50, DHSB). After primary antibody incubation, samples were washed six times in PBX (15 minutes each) at RT. Samples were then incubated in appropriate secondary antibodies (1:500, goat-anti-chicken/AF488; A11039, goat-anti-mouse/AF488, A21221, goat-anti-mouse/AF568, A11031, goat-anti-rabbit/AF568, A11011, goat-anti-rat/AF647, A21236, Thermo Fisher) an overnight at 4°C. Following secondary antibody incubation, samples underwent 6 additional 15-miniute PBS washes were mounted in VECTASHIELD anti-fade mounting medium. Samples were imaged using a Zeiss 710 LSM confocal microscope with a 20×/NA0.8 objective. Both qualitative and quantitative imaging were performed with identical laser power and gain setting within a single acquisition session to ensure consistency across samples. All image processing and quantification were conducted using FIJI (ImageJ).

#### Immunohistochemistry image analyses

Quantitation of OAMB expression in C4da neurons was conducted in FIJI using Trojan-GAL4-driven GFP expression. For each neuron, z-stack comprising 2-3 optical sections were projected into a single maximum intensity image to capture the full volume of the sensory neurons cluster. A region of interest (ROI) was defined via thresholding in CsChrimson-tdTomato channel (red), and any non-specific signal was excluded manually before mask application. Mean fluorescence intensity values for GFP and Elav were measured within the ROI. To normalize across samples, GFP signal intensities were divided by the corresponding Elav mean intensity within the same ROI.

### Two-photon live imaging

#### Sample preparation

Live imaging of C4da dendritic activity was performed in intact larvae. Larvae were cold-anesthetized and mounted directly onto a glass slide. After drying the surface briefly, larvae were gently immobilized between the slide and a 50x24 mm coverslip to restrict movement, ensuring that dorsal cuticle remained stable and oriented toward the objective lens. Any larvae exhibiting movement during imaging were excluded from further analysis.

For imaging octopaminergic neuron activity, isolated CNS from third instar larvae were prepared following the protocol described by *Zhu et al., 2024*^40^. Briefly, the CNS was dissected in cold Baines physiological solution, then transferred onto a poly-L-lysine-coated cover glass shard for stabilization.

#### Image acquisition and analysis

Functional calcium imaging was conducted using a custom-build two-photon microscope equipped with Galvo-Resonant Scanner (Cambridge technology), operated by ScanImage software (MBF Bioscience). A 40×/0.80NA water immersion objective (LUMPlanFL, Olympus) was used for image acquisition. GCaMP was excited at 920 nm using a Mai Tai ultrafast pulsed laser (Spectra-Physics). Emission signals were detected by GaAsP photomultiplier tubes (Hamamatsu) controlled by a HHMI PMT controller. Images were acquired at 30 frames per second (fps) on a single focal plane with spatial resolution of 512 x 512 pixels. A 620-nm LED (Thorlabs) was used for optogenetic stimulation. Stimuli were delivered in three repeated trials, with the following conditions: Whole larvae GCaMP6s imaging: 10 pulses (30 ms each) over 1,000 ms, repeated every 30 s. CNS GCaMP6s imaging: 100 pulses (9 ms each) over 1,000 ms, repeated every 60 s. GRAB-OA1.0 imaging: 500 pulses (9 ms each) over 5,000 ms, repeated every 60 s. Unless otherwise noted, the stimulation intensity was set to 0.1 μW/mm^2^.

To reduce background noise, acquired images were averaged to yield a time-lapse stack with a 10-fps temporal resolution. Image analysis was performed using FIJI and MATLAB. ROI was defined based on standard deviation projections across full time recording. Fluorescence intensity was quantified as ΔF/F_0_, where ΔF = F – F_0_ and F_0_ represents the mean baseline fluorescence during the 5 seconds preceding stimulus onset. Background fluorescence was subtracted prior to normalization. Responses were then averaged across the three stimulations trials per preparation. The following metrics were extracted from resulting traces: peak before post-processing to obtain the peak ΔF/F_0_, latency to peak, and signal decay time as the represented as the half-life from the peak.

### Statistics

The JMP 18 Pro software (SAS Institute Inc.) was used for statistical analysis. For continuous data types, the Shapiro-Wilk test was used to validate the normality of the dataset. If the assumption is respected, the groups were compared using a one-way ANOVA, with post-hoc Fisher’s LSD test used to determine pairwise comparison in cases of multiple comparisons. Welch’s ANOVA was used in case of severe discrepancies in n or unequal variance as asserted with Levene’s test. Continuous data that failed to meet assumptions and ordinal data led to the use of a non-parametric equivalent, namely the Kruskal-Wallis test followed by a non-parametric Fisher’s LSD test based on pairwise Mann-Whitney test, or a Mann-Whitney test for comparisons across two samples. Categorical data were analyzed using pairwise Chi-square tests, with a Bonferroni correction in cases of multiple comparisons. The type of statistical test used in each experiment is indicated in figure legends. Sample numbers are indicated in figure legends. P values are represented by asterisks: **: p < 0.01, ***: p < 0.001. For multiple comparisons, the compact letter display (CLD) is used at p < 0.01. Summary values represented in figures can be found in supplemental table 3. Exact p-values and tests can be found in supplemental table 4. Source data can be found in the supplemental material.

## Data availability statement

The original contributions presented in this study are included in the article/supplemental material, further inquiries can be directed to the corresponding author.

## Author contributions

Conceptualization, J-C.B. and T.O. Writing – Original Draft, J-C.B. and T.O, Writing – Review & Editing, J-C.B., Y.Q.Z., and T.O. Formal Analysis, J-C.B., and T.O. Performing experiments, J-C.B., Y.Q.Z., J.Z., J.T.D., J.N., and T.O., Supervision, T.O.

## Acknowledgements

Confocal images were collected at the McGill University Advanced Bio Imaging Facility (ABIF), RRID:SCR_017697. We thank for Bloomington stock center for providing the fly stocks. We thank for V. Jayaraman, G. Rubin, J. Simpson, R. S. Stowers and D. Tracy to share their fly stocks. We thank Marta Zlatic and members of Ohyama lab for critical comments on the manuscript.

