## supplemental information for "A positive feedback loop between sensory and octopaminergic neurons underlies nociceptive plasticity in *Drosophila* larvae"

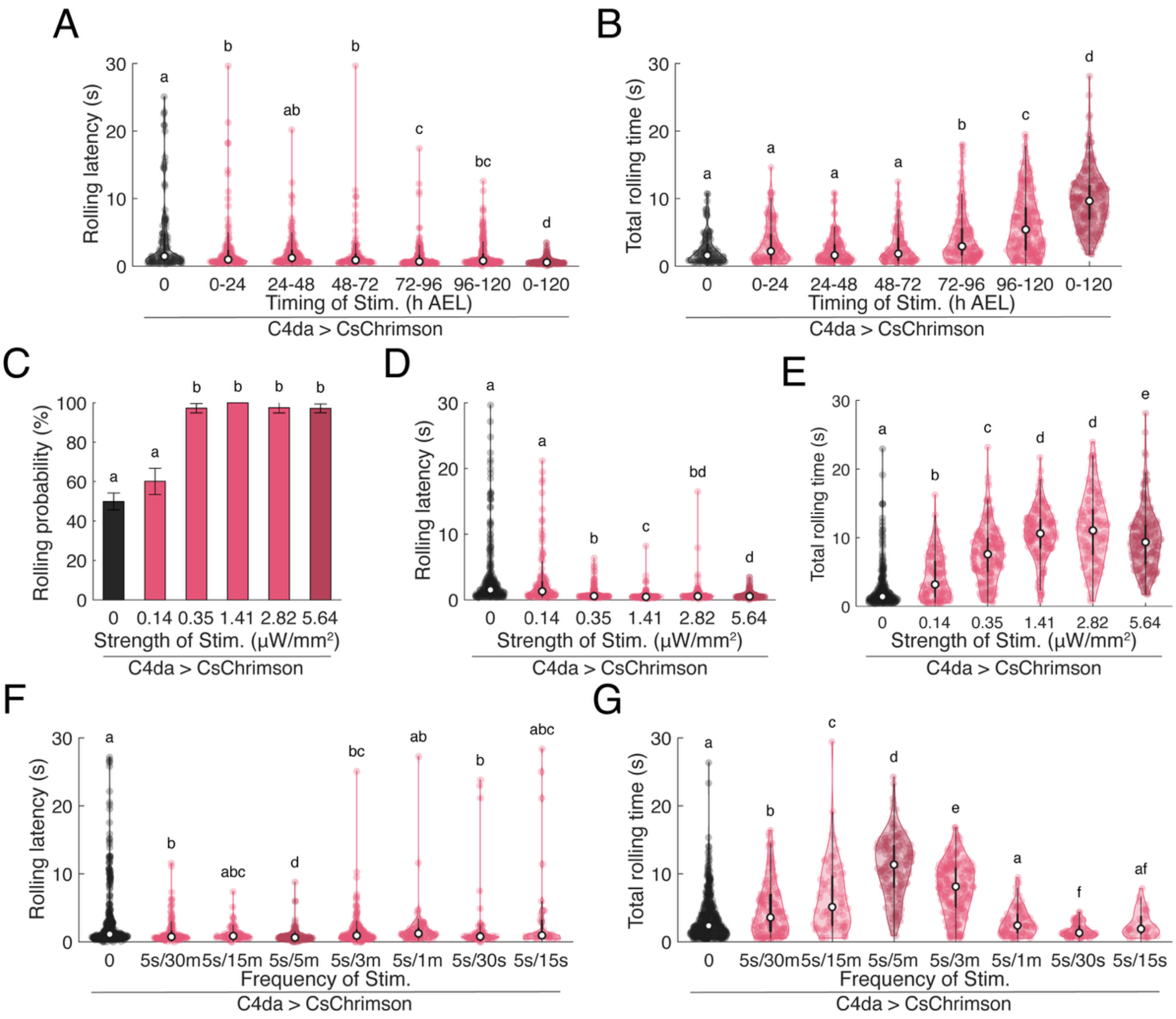

**Supplemental figure 1. Key features of developmental stimulations differentially affect rolling behaviour.**

(A-B) Developmental stage has an incremental effect on sensitization, with most potent effects with developmental stimulation of late-third instar. (A) Violin plot of rolling latency during optogenetic stimulation (n = 106, 125 138, 113, 165, 229, 254). Kruskall-Wallis followed by pairwise Mann-Whitney test, CLD denotes $p < 0.01$. (B) Violin plot of total time each larvae spent rolling during optogenetic stimulations (n = 106, 125 138, 113, 165, 229, 254). Kruskall-Wallis followed by pairwise Mann-Whitney test; CLD denotes $p < 0.01$.

(C-E) Stimulation intensity increases sensitization, but the effect rapidly saturates. (C) Rolling probability during optogenetic C4da stimulation. Error bars: 95% confidence interval (n = 517, 208, 185, 160, 124, 214). Chi-square test with Bonferroni correction; CLD denotes $p < 0.01$. (D) Violin plot of rolling latency during optogenetic stimulation (n = 258, 125, 180, 160, 121, 208). Kruskall-Wallis followed by pairwise Mann-Whitney test, CLD denotes $p < 0.01$. (E) Violin plot of total time each larvae spent rolling during optogenetic stimulations (n = 258, 125, 180, 160, 121, 208). Kruskall-Wallis followed by pairwise Mann-Whitney test; CLD denotes $p < 0.01$.

(F-G) Stimulation frequency dictates directionality of behavioural changes. (F) Violin plot of rolling latency during optogenetic stimulation (n = 407, 147, 75, 172, 215, 92, 67, 35). Kruskall-Wallis followed by pairwise Mann-Whitney test, CLD denotes $p < 0.01$. (G) Violin plot of total time each larvae spent rolling during optogenetic stimulations (n = 106, 125 138, 113, 165, 229, 254). Kruskall-Wallis followed by pairwise Mann-Whitney test; CLD denotes $p < 0.01$.

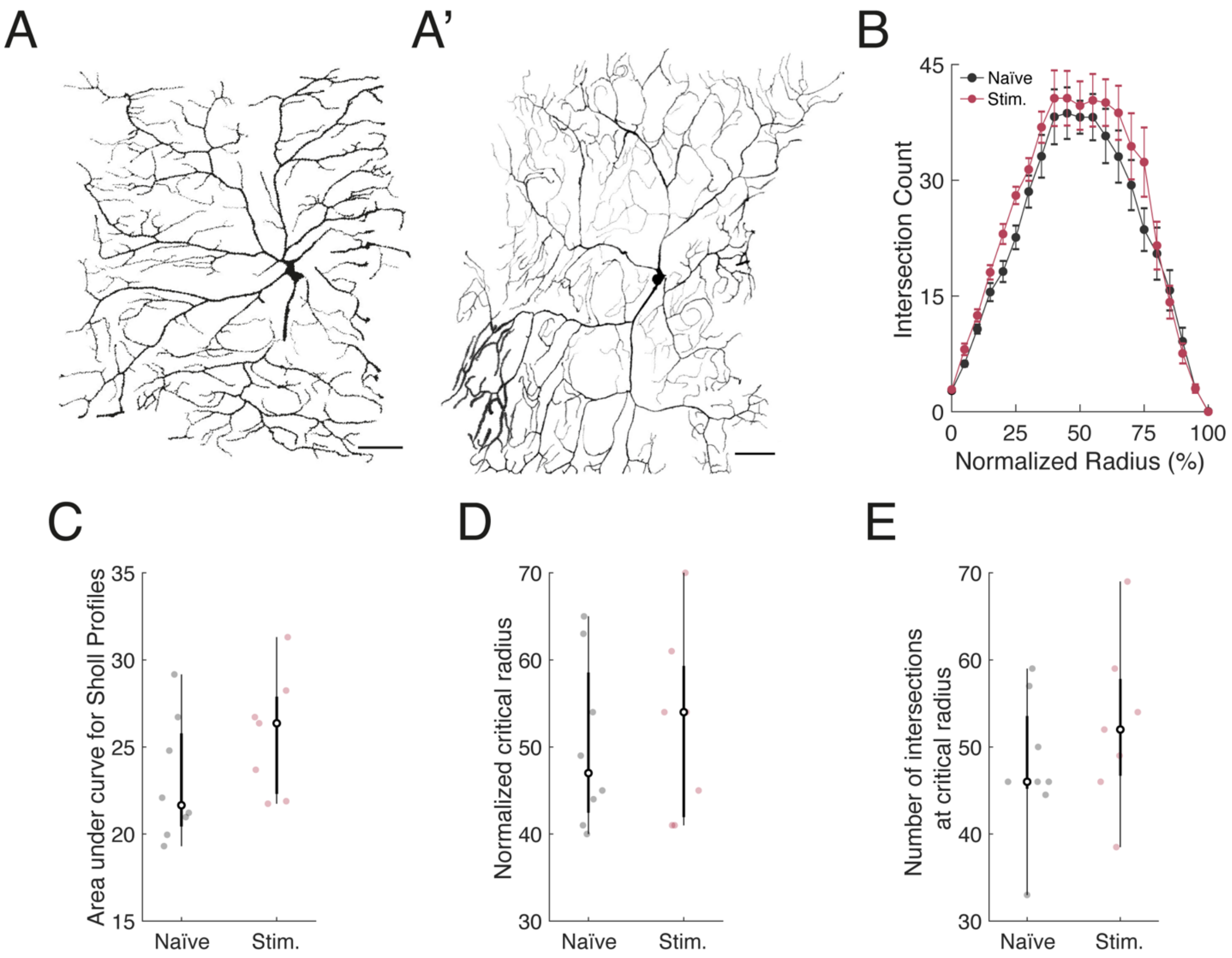

**Supplemental figure 2. Developmental stimulations do not affect C4da dendrite morphology.**

(A-A') Representative dendritic arbour from naïve and developmentally stimulated larvae. Scale bar = 50 µm. (A) Naïve. (A') Stimulated.

(B) Average Sholl profiles for naïve (n=7) and stimulated larvae (n=8). Profiles were normalized to control for inter-individual variance in larval size.

(C-E) Boxplots representing different core features of the normalized Sholl profiles in naïve (n=7) and stimulated larvae (n=8). (C) Area under the curve. (D) Critical radius. (E) Number of intersection at critical radius.

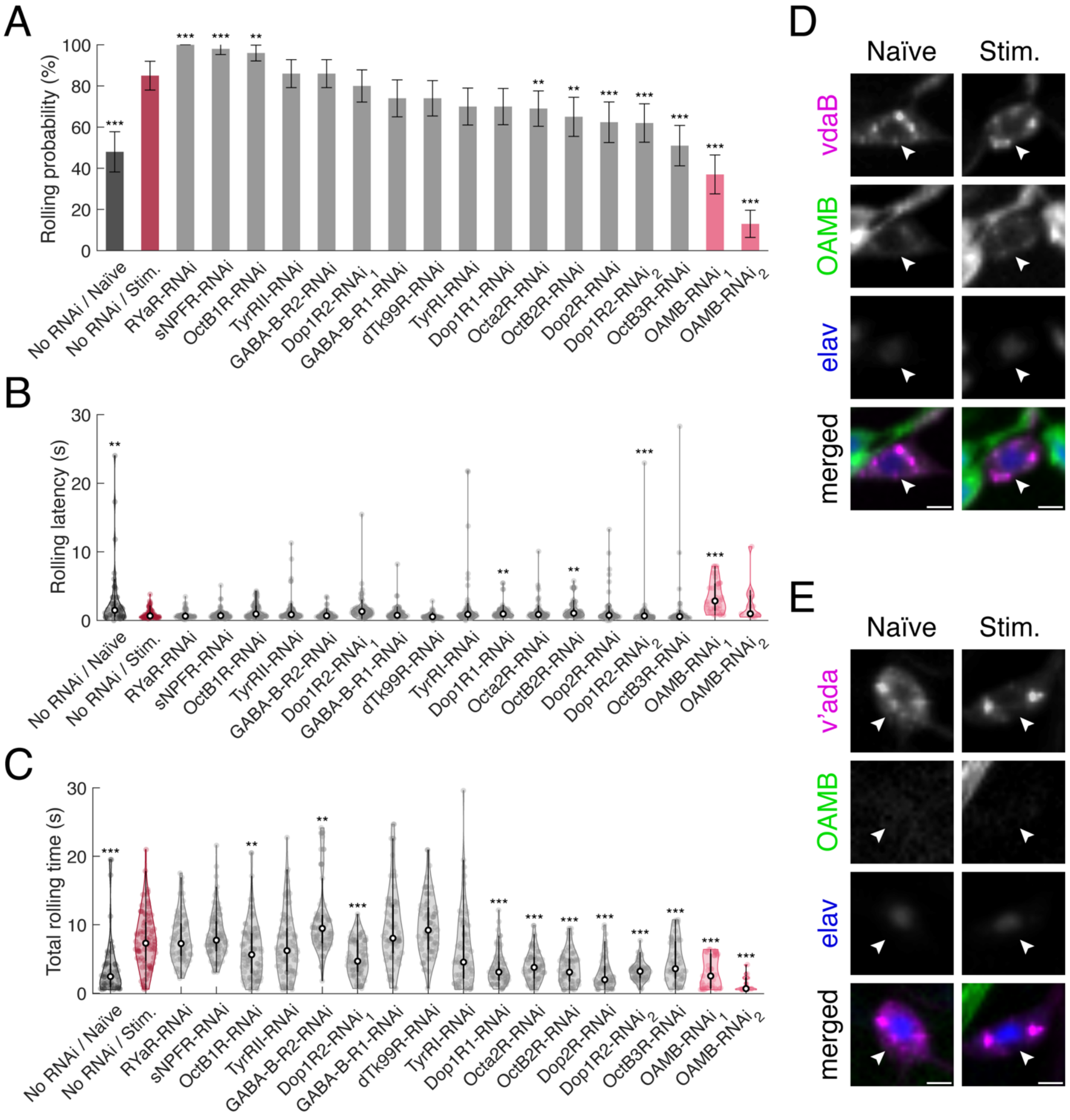

**Supplemental figure 3. The octopamine receptor OAMB is required for experience-dependent sensitization to noxious stimuli.**

(A-C) An RNAi screening identifies OAMB in C4da neurons as critical for sensitization following noxious experience. (A) Rolling probability during optogenetic C4da stimulation. Error bars: 95% confidence interval (n = 100, 100, 100, 100, 100, 93, 100, 100, 100, 100, 100, 100, 100, 100, 100, 70, 100, 100, 100). Chi-square test; statistical marks are relative to the No RNAi / Stim. group. (B) Violin plot of rolling latency during optogenetic stimulation (n = 48, 85, 70, 80, 62, 58, 74, 74, 86, 37, 13, 69, 96, 65, 51, 70, 98, 70, 86). Kruskall-Wallis followed by pairwise a Steel-Dwass test with control (No RNAi / Stim.) (C) Violin plot of total time each larvae spent rolling during optogenetic stimulations (n = 48, 85, 70, 80, 62, 58, 74, 74, 86, 37, 13, 69, 96, 65, 51, 70, 98, 70, 86). Kruskall-Wallis followed by pairwise a Steel-Dwass test with control (No RNAi / Stim.).

(D-E) OAMB expression varies with neuronal identity. (D) Representative images mean OAMB expression in vdaB neurons in naïve and stimulated larvae. Scale bar = 10 µm. (E) Representative images mean OAMB expression in v'ada neurons in naïve and stimulated larvae. Scale bar = 10 µm

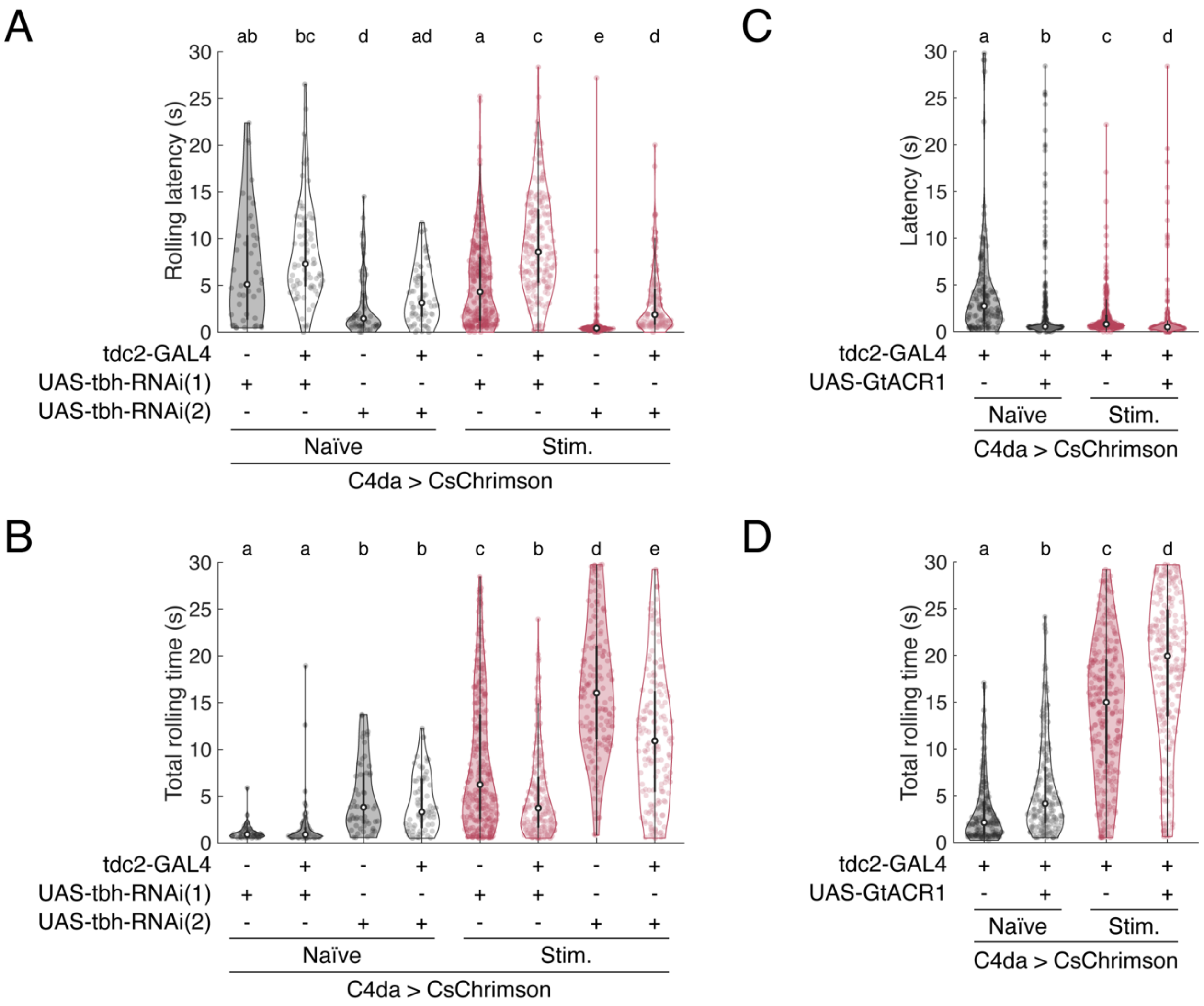

**Supplemental figure 4. Octopaminergic signalling from tdc2+ neurons affects larval rolling behaviour following developmental stimulations.**

(A-B) RNAi knockdown of the rate-limiting enzyme tbh affects rolling latency and total duration. (A) Violin plot of rolling latency during optogenetic stimulation (n =44, 83, 55, 70, 292, 226, 163, 167). Kruskall-Wallis followed by pairwise Mann-Whitney test, CLD denotes $p < 0.01$. (B) Violin plot of total time each larvae spent rolling during optogenetic stimulations (n =44, 83, 55, 70, 292, 226, 163, 167). Kruskall-Wallis followed by pairwise Mann-Whitney test; CLD denotes $p < 0.01$.

(C-D) Optogenetic silencing during developmental stimulations affects rolling latency but not total duration (C) Violin plot of rolling latency during optogenetic stimulation (n =263, 144, 265, 217). Kruskall-Wallis followed by pairwise Mann-Whitney test, CLD denotes $p < 0.01$. (D) Violin plot of total time each larvae spent rolling during optogenetic stimulations (n =263, 144, 265, 217). Kruskall-Wallis followed by pairwise Mann-Whitney test; CLD denotes $p < 0.01$.

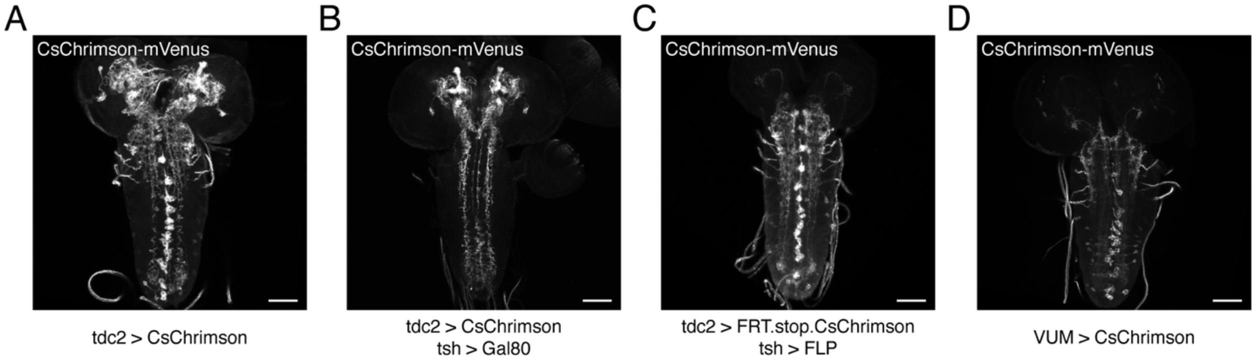

**Supplemental figure 5. Combinations of genetic reagents allow for isolation different tdc2+ neuron clusters.**

(A-D) Example images of different techniques used to drive CsChrimson in different tdc2+ neuron clusters. Scale bar = 50 µm. (A) Expression of UAS-CsChrimson driven by tdc2-GAL4. (B) Expression of UAS-CsChrimson driven by tdc2-GAL4 combined to GAL80 expression driven by tsh-LexA, allowing for isolation of octopaminergic neurons of the brain. (C) Expression of UAS-FRT.Stop-CsChrimson driven by tdc2-GAL4 combined to FLP expression driven by tsh-LexA, allowing for isolation of octopaminergic neurons of the ventral nerve chord. (D) Expression of UAS-CsChrimson driven by 46B08-GAL4, allowing for isolation of the VUMs.

Supplemental Table 1. Number of hemisegments in which C4da neurons from synapses with tdc2+ neurons

| | Presence of GRASP Signal | | | |
|---|---|---|---|---|
| | Naïve | | Stim. | |
| Segment | Left | Right | Left | Right |
| T1 | 0 | 1 | 5 | 5 |
| T2 | 10 | 11 | 6 | 5 |
| T3 | 8 | 6 | 3 | 4 |
| A1 | 4 | 3 | 5 | 5 |
| A2 | 8 | 10 | 11 | 13 |
| A3 | 12 | 11 | 11 | 12 |
| A4 | 8 | 9 | 11 | 11 |
| A5 | 6 | 6 | 12 | 13 |
| A6 | 9 | 4 | 14 | 13 |
| A7 | 6 | 10 | 14 | 14 |
| A8/9 | 11 | 12 | 14 | 14 |
| n | 14 | | 14 | |

Supplemental Table 2. The flies carrying the following constructs were used for experiments and were represented in figures as follows.

| Figure | Label | Genotype |
|---|---|---|
| Fig1 A-G, SuppFig1 F-G, Fig2 A-F, Fig5 A-D | C4da > CsChrimson | UAS-CsChrimson-mVenus/ w; ppk1.9-GAL4 / +; + |
| Fig2 G-I, Fig5 E-F | C4da > GCaMP6s, CsChrimson | w; ppk-LexA / ppk1.9-GAL4; UAS-GCaMP6s, LexAop-Chrimson-tdTomato /+ |
| Fig3 A-C | No RNAi | w; ppk1.9-GAL4 / UAS-dcr2; UAS-CsChrimson-mVenus/P{CaryP}attP2 |
| Fig3 A-C | OAMB-RNAi(1) | w; ppk1.9-GAL4 / UAS-dcr2; UAS-CsChrimson-mVenus/TRIP.JF01732 |
| Fig3 A-C | OAMB-RNAi(2) | w; ppk1.9-GAL4 / UAS-dcr2; UAS-CsChrimson-mVenus/TRIP.JF01673 |
| Fig3 E-F | No RNAi | 20xUAS-Syn21-opGCaMP6s,10xUAS-Syn21-CsChrimson88::tdTomato / w; UAS-dcr2 / ppk-GAL4;+ |
| Fig3 E-F | OAMB-RNAi | 20xUAS-Syn21-opGCaMP6s,10xUAS-Syn21-CsChrimson88::tdTomato / w; UAS-dcr2 / ppk-GAL4;TRIP.JF01673 / + |
| Fig3 G-H, SuppFig3 D-E | | w; 27H06-LexA / + ; LexAop-Chrimson-tdTomato / Mi{Trojan-GAL4.1}Oamb[MI12417-TG4.1], UAS-myr::GFP |
| Fig4 A, SuppFig4 A-B | C4da > CsChrimson, tbh-RNAi(1) > - | LexAop-Chrimson-tdTomato / w ; ppk-LexA, UAS-dcr2 / + ; TRiP.JF02746 / P{CaryP}attP2 |
| Fig4 A, SuppFig4 A-B | C4da > CsChrimson, tbh-RNAi(1) > tdc2 | LexAop-Chrimson-tdTomato / w; ppk-LexA, UAS-dcr2 / +; TRiP.JF02746 / tdc2-GAL4.S |
| Fig4 A, SuppFig4 A-B | C4da > CsChrimson, tbh-RNAi(2) > - | ppk-LexA / HMS05829 ; LexAop-Chrimson-tdTomato / P{CaryP}attP2 |
| Fig4 A, SuppFig4 A-B | C4da > CsChrimson, tbh-RNAi(2) > tdc2 | ppk-LexA / HMS05829 ; LexAop-Chrimson-tdTomato / tdc2-GAL4.S |
| Fig4 C, SuppFig4 C-D | C4da > CsChrimson, - > tdc2 | LexAop-Chrimson-tdTomato; ppk-lexA; UAS-GtACR1 / P{CaryP}attP2 |
| Fig4 C, SuppFig4 C-D | C4da > CsChrimson, GtACR1 > tdc2 | LexAop-Chrimson-tdTomato; ppk-lexA; UAS-GtACR1 / tdc2-GAL4.S |
| Fig6 A | | 13xLexAop-mCD8::GFP (su(Hw)attP8),10XUAS-IVS-mCD8::RFP (attP18) / w; ppk1.9-GAL4 / tdc2-LexA |

| | | |
|---|---|---|
| Fig6 C-D | No GAL4 | UAS-Chrimson-mVenus (attp18) / w; +; tdc2-GAL4.S / P{CaryP}attP2 |
| Fig6 C-D, SuppFig6 A | tdc2-GAL4 | UAS-Chrimson-mVenus (attp18) / w; +; tdc2-GAL4.S /+ |
| Fig6 C-D, SuppFig6 B | tsh-GAL80 | UAS-Chrimson-mVenus (attp18) / w; tsh-LexA,LexAop-GAL80 / +; tdc2-GAL4.S /+ |
| Fig6 C-D, SuppFig6 C | tsh-FLP | UAS-FRT.Stop-Chrimson-mVenus (attp18) / w ;tsh-LexA,LexAop-FLP / +; tdc2-GAL4.S / + |
| Fig6 C-D, SuppFig6 D | VUM-GAL4 | UAS-Chrimson-mVenus (attp18) / w; +; GMR46B08-GAL4 /+ |
| Fig6 E | | LexAop-nsyb::GFP1-10, UAS-CD4::GFP11 / ppk-LexA; LexAop-Chrimson-tdTomato / tdc2-GAL4.S |
| Fig6 F-G | C4da > CsChrimson ; tdc2 > GCaMP6s | w; ppk-LexA / +; UAS-GCaMP6s, LexAop-Chrimson-tdTomato / tdc2-GAL4.S |
| Fig6 H-I | tdc2 > CsChrimson ; C4da > GRAB-OA1.0 | w; UAS-GRAB(OA1.0), ppk1.9-GAL4 / tdc2-LexA; LexAop-Chrimson-tdTomato / + |
| SuppFig2 A-E | | UAS-CsChrimson-mVenus / w; ppk-CD4-tdTomato, ppk1.9-GAL4 / +; + |

Supplemental Table 3. Summary values for the data represented in figures.

| Fig1-C | | | |
|---|---|---|---|
| GroupID | n | Rolling probability (%) | 95% CI |
| Naïve | 327 | 42.20 | 5.35 |
| Stim. | 207 | 98.07 | 1.87 |
| Fig1-D | | | |
| GroupID | n | Latency (s) | SEM |
| Naïve | 138 | 4.30 | 0.49 |
| Stim. | 203 | 0.67 | 0.03 |
| Fig1-E | | | |
| GroupID | n | Total rolling time (s) | SEM |
| Naïve | 138 | 2.73 | 0.30 |
| Stim. | 203 | 10.11 | 0.31 |
| Fig1-F | | | |
| GroupID | n | Rolling probability (%) | 95% CI |
| Naïve | 237 | 44.73 | 6.33 |
| 0-24h | 234 | 53.42 | 6.39 |
| 24-48h | 267 | 51.69 | 5.99 |
| 48-72h | 187 | 60.43 | 7.01 |
| 72-96h | 211 | 78.20 | 5.57 |
| 96-120h | 283 | 80.82 | 4.59 |
| 0-120h | 260 | 97.69 | 1.83 |
| Fig1-G | | | |
| GroupID | n | Rolling probability (%) | 95% CI |
| Naïve | 818 | 49.76 | 3.43 |
| 5s/30m | 277 | 53.07 | 5.88 |
| 5s/15m | 100 | 75.00 | 8.49 |
| 5s/5m | 174 | 98.85 | 1.58 |
| 5s/3m | 219 | 98.17 | 1.78 |
| 5s/m | 194 | 47.42 | 7.03 |
| 5s/30s | 190 | 35.26 | 6.79 |
| 5s/15s | 168 | 20.83 | 6.14 |
| Fig2-B | | | |
| Group-Behaviour | n | Behaviour Probability (%) | 95% CI |
| Naïve-Crawl | 5 | 3.42 | 2.95 |
| Naïve-Stop | 37 | 25.34 | 7.06 |
| Naïve-Cast | 12 | 8.22 | 4.46 |
| Naïve-C-shape | 36 | 24.66 | 6.99 |

| | | | |
|---|---|---|---|
| Naïve-Roll | 56 | 38.36 | 7.89 |
| Stim.-Crawl | 2 | 1.18 | 1.63 |
| Stim.-Stop | 15 | 8.88 | 4.29 |
| Stim.-Cast | 9 | 5.33 | 3.39 |
| Stim.-C-shape | 30 | 17.75 | 5.76 |
| Stim.-Roll | 113 | 66.86 | 7.10 |
| Fig2-C | | | |
| Group | n | Rolling probability (%) | 95% CI |
| Naïve | 343 | 79.30 | 4.29 |
| Stim | 350 | 92.86 | 2.70 |
| Group | n | Latency (s) | SEM |
| Naïve | 343 | 4.30 | 0.15 |
| Stim | 350 | 3.27 | 0.12 |
| Fig2-D | | | |
| Group | n | Rolling probability (%) | 95% CI |
| Naïve-20mN | 50 | 0.00 | 0.00 |
| Naïve-40mN | 50 | 10.00 | 8.32 |
| Naïve-60mN | 50 | 20.00 | 11.09 |
| Naïve-80mN | 50 | 48.00 | 13.85 |
| Naïve-100mN | 50 | 58.00 | 13.68 |
| Naïve-120mN | 50 | 74.00 | 12.16 |
| Naïve-150mN | 50 | 78.00 | 11.48 |
| Stim.-20mN | 50 | 14.00 | 9.62 |
| Stim.-40mN | 50 | 26.00 | 12.16 |
| Stim.-60mN | 50 | 64.00 | 13.30 |
| Stim.-80mN | 50 | 82.00 | 10.65 |
| Stim.-100mN | 50 | 96.00 | 5.43 |
| Stim.-120mN | 50 | 100.00 | 0.00 |
| Stim.-150mN | 50 | 100.00 | 0.00 |
| Fig2-E | | | |
| Group | n | Rolling probability (%) | 95% CI |
| Naïve-0.1% | 50 | 0.00 | 0.00 |
| Naïve-0.5% | 50 | 0.00 | 0.00 |
| Naïve-1% | 50 | 40.00 | 13.58 |
| Naïve-2.5% | 50 | 68.00 | 12.93 |
| Naïve-5% | 50 | 76.00 | 11.84 |
| Naïve-7.5% | 50 | 84.00 | 10.16 |
| Naïve-10% | 50 | 94.00 | 6.58 |

| | | | |
|---|---|---|---|
| Stim.-0.1% | 50 | 2.00 | 3.88 |
| Stim.-0.5% | 50 | 46.00 | 13.81 |
| Stim.-1% | 50 | 64.00 | 13.30 |
| Stim.-2.5% | 50 | 86.00 | 9.62 |
| Stim.-5% | 50 | 100.00 | 0.00 |
| Stim.-7.5% | 50 | 100.00 | 0.00 |
| Stim.-10% | 50 | 98.00 | 3.88 |
| Fig2-F | | | |
| Group | n | Rolling probability (%) | 95% CI |
| Naive-0.01 | 110 | 3.64 | 3.50 |
| Naive-0.03 | 189 | 2.65 | 2.29 |
| Naive-0.04 | 142 | 11.27 | 5.20 |
| Naive-0.07 | 92 | 26.09 | 8.97 |
| Naive-0.1 | 305 | 37.38 | 5.43 |
| Stim.-0.01 | 116 | 3.45 | 3.32 |
| Stim.-0.03 | 189 | 9.52 | 4.19 |
| Stim.-0.04 | 117 | 25.64 | 7.91 |
| Stim.-0.07 | 184 | 38.04 | 7.02 |
| Stim.-0.1 | 129 | 68.99 | 7.98 |
| Fig2-I | | | |
| Group | n | Maximum dF/F0 | SEM |
| Naïve-0.1 | 12 | 1.51 | 0.19 |
| Naïve-0.3 | 13 | 2.18 | 0.29 |
| Naïve-0.5 | 12 | 2.85 | 0.53 |
| Naïve-0.9 | 12 | 2.97 | 0.45 |
| Naïve-1.4 | 11 | 3.53 | 0.82 |
| Stim.-0.1 | 10 | 4.72 | 1.03 |
| Stim.-0.3 | 12 | 5.19 | 1.05 |
| Stim.-0.5 | 10 | 6.15 | 0.86 |
| Stim.-0.9 | 11 | 7.19 | 1.08 |
| Stim.-1.4 | 10 | 7.34 | 1.19 |
| Fig3-A | | | |
| Group | n | Rolling probability (%) | 95% CI |
| No RNAi / Naïve | 589 | 33.11 | 3.80 |
| OAMB-RNAi_1 / Naïve | 283 | 20.49 | 4.70 |
| OAMB-RNAI_2 / Naïve | 270 | 31.85 | 5.56 |
| No RNAi / Stim. | 284 | 94.57 | 2.64 |
| OAMB-RNAi_1 / Stim. | 278 | 35.97 | 5.64 |
| OAMB-RNAI_2 / Stim. | 414 | 11.84 | 3.11 |

| | | | |
|---|---|---|---|
| Fig3-B | | | |
| GroupID | n | Latency (s) | SEM |
| No RNAi / Naïve | 194 | 2.70 | 0.27 |
| OAMB-RNAi_1 / Naïve | 58 | 3.94 | 0.67 |
| OAMB-RNAI_2 / Naïve | 86 | 4.23 | 0.58 |
| No RNAi / Stim. | 268 | 0.87 | 0.04 |
| OAMB-RNAi_1 / Stim. | 100 | 2.85 | 0.33 |
| OAMB-RNAI_2 / Stim. | 49 | 5.50 | 1.20 |
| Fig3-C | | | |
| GroupID | n | Total rolling time (s) | SEM |
| No RNAi / Naïve | 195 | 2.50 | 0.20 |
| OAMB-RNAi_1 / Naïve | 58 | 1.87 | 0.24 |
| OAMB-RNAI_2 / Naïve | 86 | 1.97 | 0.20 |
| No RNAi / Stim. | 268 | 6.76 | 0.18 |
| OAMB-RNAi_1 / Stim. | 100 | 3.05 | 0.22 |
| OAMB-RNAI_2 / Stim. | 49 | 1.43 | 0.19 |
| Fig3-F | | | |
| GroupID | n | Normalized fluorescence | SEM |
| No RNAi / Naïve | 10 | 0.63 | 0.10 |
| No RNAi / Stim. | 9 | 1.12 | 0.07 |
| OAMB-RNAi / Naïve | 10 | 0.62 | 0.11 |
| OAMB-RNAi / Stim. | 9 | 0.52 | 0.08 |
| Fig3-H | | | |
| GroupID | n | Normalized fluorescence | SEM |
| ddaC / Naïve | 6 | 0.12 | 0.03 |
| ddaC / Stim. | 8 | 0.40 | 0.03 |
| v'ada / Naïve | 6 | 0.04 | 0.03 |
| v'ada / Stim. | 8 | 0.06 | 0.01 |
| vdaB / Naïve | 6 | 0.63 | 0.13 |
| vdaB / Stim. | 8 | 0.76 | 0.09 |
| Fig4-A | | | |
| GroupID | n | Rolling probability (%) | 95% CI |
| tbh-RNAi_1>- / Dev. Stim. | 351 | 83.19 | 3.91 |
| tbh-RNAi_1>- / No Dev. Stim. | 334 | 13.17 | 3.63 |
| tbh-RNAi_1>tdc2 / Dev. Stim. | 343 | 65.89 | 5.02 |
| tbh-RNAi_1>tdc2 / No Dev. Stim. | 579 | 14.34 | 2.85 |
| tbh-RNAi_2>- / Dev. Stim. | 211 | 77.73 | 5.61 |
| tbh-RNAi_2>- / No Dev. Stim. | 190 | 28.95 | 6.45 |

| | | | |
|---|---|---|---|
| tbh-RNAi_2>tdc2 / Dev. Stim. | 285 | 58.60 | 5.72 |
| tbh-RNAi_2>tdc2 / No Dev. Stim. | 300 | 23.33 | 4.79 |
| Fig4-C | | | |
| GroupID | n | Rolling probability (%) | 95% CI |
| attp2 / Dev. Stim. | 306 | 85.95 | 3.89 |
| attp2 / No Dev. Stim. | 391 | 36.83 | 4.78 |
| tdc2-GAL4 / Dev. Stim. | 455 | 58.24 | 4.53 |
| tdc2-GAL4 / No Dev. Stim. | 483 | 44.93 | 4.44 |
| Fig5-B | | | |
| GroupID | n | Rolling probability (%) | 95% CI |
| 0mM | 194 | 48.97 | 7.03 |
| 5mM | 217 | 69.12 | 6.15 |
| 25mM | 209 | 72.25 | 6.07 |
| 50mM | 174 | 77.01 | 6.25 |
| Fig5-C | | | |
| GroupID | n | Rolling probability (%) | 95% CI |
| 0mM-Crawl | 100 | 8.00 | 5.32 |
| 0mM-Stop | 100 | 32.00 | 9.14 |
| 0mM-Turn | 100 | 6.00 | 4.65 |
| 0mM-C-shape | 100 | 23.00 | 8.25 |
| 0mM-Roll | 100 | 31.00 | 9.06 |
| 5mM-Crawl | 100 | 5.00 | 4.27 |
| 5mM-Stop | 100 | 19.00 | 7.69 |
| 5mM-Turn | 100 | 5.00 | 4.27 |
| 5mM-C-shape | 100 | 19.00 | 7.69 |
| 5mM-Roll | 100 | 52.00 | 9.79 |
| Fig5-D | | | |
| Group | n | Rolling probability (%) | 95% CI |
| 0mM | 100 | 92.00 | 5.32 |
| 5mM | 100 | 98.00 | 2.74 |
| Group | n | Latency (s) | SEM |
| 0mM | 100 | 3.16 | 0.18 |
| 5mM | 100 | 2.53 | 0.14 |
| Fig5-F | | | |
| Group | n | Maximum dF/F0 | SEM |
| 5mM OA-0.1 | 10 | 2.13 | 0.27 |
| Untreated-0.1 | 10 | 1.42 | 0.20 |

| | | | |
|---|---|---|---|
| 5mM OA-0.5 | 12 | 3.52 | 0.37 |
| Untreated-0.5 | 10 | 2.33 | 0.19 |
| 5mM OA-1.4 | 10 | 4.46 | 0.67 |
| Untreated-1.4 | 10 | 2.95 | 0.35 |
| Fig6-C | | | |
| Group | n | Behaviour Probability (%) | 95% CI |
| attp2-Naive-Crawl | 205 | 3.90 | 2.65 |
| attp2-Naive-Stop | 205 | 30.73 | 6.32 |
| attp2-Naive-Cast | 205 | 7.81 | 3.67 |
| attp2-Naive-Cshape | 205 | 20.49 | 5.53 |
| attp2-Naive-Roll | 205 | 37.07 | 6.61 |
| attp2-Stim-Crawl | 198 | 3.03 | 2.39 |
| attp2-Stim-Stop | 198 | 29.29 | 6.34 |
| attp2-Stim-Cast | 198 | 8.59 | 3.90 |
| attp2-Stim-Cshape | 198 | 21.21 | 5.69 |
| attp2-Stim-Roll | 198 | 37.88 | 6.76 |
| tdc2-tsh-gal80-Naive-Crawl | 100 | 8.00 | 5.32 |
| tdc2-tsh-gal80-Naive-Stop | 100 | 17.00 | 7.36 |
| tdc2-tsh-gal80-Naive-Cast | 100 | 8.00 | 5.32 |
| tdc2-tsh-gal80-Naive-Cshape | 100 | 26.00 | 8.60 |
| tdc2-tsh-gal80-Naive-Roll | 100 | 41.00 | 9.64 |
| tdc2-tsh-gal80-Stim-Crawl | 100 | 8.00 | 5.32 |
| tdc2-tsh-gal80-Stim-Stop | 100 | 19.00 | 7.69 |
| tdc2-tsh-gal80-Stim-Cast | 100 | 7.00 | 5.00 |
| tdc2-tsh-gal80-Stim-Cshape | 100 | 27.00 | 8.70 |
| tdc2-tsh-gal80-Stim-Roll | 100 | 39.00 | 9.56 |
| tdc2-Naive-Crawl | 100 | 8.00 | 5.32 |
| tdc2-Naive-Stop | 100 | 24.00 | 8.37 |
| tdc2-Naive-Cast | 100 | 10.00 | 5.88 |
| tdc2-Naive-Cshape | 100 | 25.00 | 8.49 |
| tdc2-Naive-Roll | 100 | 33.00 | 9.22 |
| tdc2-Stim-Crawl | 100 | 4.00 | 3.84 |
| tdc2-Stim-Stop | 100 | 16.00 | 7.19 |
| tdc2-Stim-Cast | 100 | 5.00 | 4.27 |
| tdc2-Stim-Cshape | 100 | 23.00 | 8.25 |
| tdc2-Stim-Roll | 100 | 52.00 | 9.79 |
| tdc2-tsh-FLP-Naive-Crawl | 100 | 13.00 | 6.59 |
| tdc2-tsh-FLP-Naive-Stop | 100 | 13.00 | 6.59 |

| | | | |
|---|---|---|---|
| tdc2-tsh-FLP-Naive-Cast | 100 | 12.00 | 6.37 |
| tdc2-tsh-FLP-Naive-Cshape | 100 | 38.00 | 9.51 |
| tdc2-tsh-FLP-Naive-Roll | 100 | 34.00 | 9.28 |
| tdc2-tsh-FLP-Stim-Crawl | 100 | 5.00 | 4.27 |
| tdc2-tsh-FLP-Stim-Stop | 100 | 15.00 | 7.00 |
| tdc2-tsh-FLP-Stim-Cast | 100 | 5.00 | 4.27 |
| tdc2-tsh-FLP-Stim-Cshape | 100 | 21.00 | 7.98 |
| tdc2-tsh-FLP-Stim-Roll | 100 | 54.00 | 9.77 |
| VUM-Naive-Crawl | 100 | 13.00 | 6.59 |
| VUM-Naive-Stop | 100 | 16.00 | 7.19 |
| VUM-Naive-Cast | 100 | 15.00 | 7.00 |
| VUM-Naive-Cshape | 100 | 22.00 | 8.12 |
| VUM-Naive-Roll | 100 | 34.00 | 9.28 |
| VUM-Stim-Crawl | 100 | 4.00 | 3.84 |
| VUM-Stim-Stop | 100 | 14.00 | 6.80 |
| VUM-Stim-Cast | 100 | 6.00 | 4.65 |
| VUM-Stim-Cshape | 100 | 24.00 | 8.37 |
| VUM-Stim-Roll | 100 | 52.00 | 9.79 |
| Fig6-D | | | |
| Group | n | Latency (s) | SEM |
| attp2-Naïve | 113 | 5.21 | 0.33 |
| attp2-Stim. | 112 | 4.68 | 0.30 |
| tdc2-tsh-gal80-Naïve | 100 | 4.69 | 0.29 |
| tdc2-tsh-gal80-Stim. | 100 | 4.72 | 0.28 |
| tdc2-Naïve | 100 | 4.74 | 0.30 |
| tdc2-Stim. | 100 | 3.25 | 0.21 |
| tdc2-tsh-FLP-Naïve | 100 | 4.50 | 0.30 |
| tdc2-tsh-FLP-Stim. | 100 | 2.97 | 0.18 |
| VUM-Naïve | 100 | 4.34 | 0.29 |
| VUM-Stim | 100 | 3.07 | 0.23 |
| Fig6-G | | | |
| Group | n | Maximum dF/F0 | SEM |
| Naïve | 10 | 0.77 | 0.12 |
| Stim | 13 | 2.67 | 0.36 |
| Fig6-I | | | |
| Group | n | Maximum dF/F0 | SEM |
| Naïve | 10 | 0.77 | 0.12 |
| Stim | 13 | 2.67 | 0.36 |

| SuppFig1-A | | | |
|---|---|---|---|
| GroupID | n | Latency (s) | SEM |
| Naïve | 106 | 4.18 | 0.56 |
| 0-24h | 125 | 2.53 | 0.39 |
| 24-48h | 138 | 2.11 | 0.22 |
| 48-72h | 113 | 1.89 | 0.31 |
| 72-96h | 165 | 1.42 | 0.16 |
| 96-120h | 229 | 1.65 | 0.13 |
| 0-120h | 254 | 0.72 | 0.03 |
| SuppFig1-B | | | |
| GroupID | n | Total rolling time (s) | SEM |
| Naïve | 106 | 2.44 | 0.23 |
| 0-24h | 125 | 3.31 | 0.27 |
| 24-48h | 138 | 2.41 | 0.18 |
| 48-72h | 113 | 2.76 | 0.23 |
| 72-96h | 165 | 4.26 | 0.29 |
| 96-120h | 229 | 6.04 | 0.29 |
| 0-120h | 254 | 9.63 | 0.26 |
| SuppFig1-C | | | |
| GroupID | n | Rolling probability (%) | 95% CI |
| Naïve | 517 | 49.90 | 4.31 |
| 0.14 | 208 | 60.10 | 6.66 |
| 0.35 | 185 | 97.30 | 2.34 |
| 1.41 | 160 | 100.00 | 0.00 |
| 2.82 | 124 | 97.58 | 2.70 |
| 5.64 | 214 | 97.20 | 2.21 |
| SuppFig1-D | | | |
| GroupID | n | Latency (s) | SEM |
| Naïve | 258 | 3.36 | 0.31 |
| 0.14 | 125 | 2.98 | 0.38 |
| 0.35 | 180 | 0.94 | 0.07 |
| 1.41 | 160 | 0.64 | 0.06 |
| 2.82 | 121 | 0.92 | 0.15 |
| 5.64 | 208 | 0.71 | 0.04 |
| SuppFig1-E | | | |
| GroupID | n | Total rolling time (s) | SEM |
| Naïve | 258 | 2.64 | 0.20 |
| 0.14 | 125 | 4.17 | 0.29 |

| | | | |
|---|---|---|---|
| 0.35 | 180 | 7.63 | 0.30 |
| 1.41 | 160 | 10.49 | 0.29 |
| 2.82 | 121 | 11.14 | 0.45 |
| 5.64 | 208 | 9.50 | 0.30 |
| SuppFig1-F | | | |
| GroupID | n | Latency (s) | SEM |
| Naïve | 407 | 2.99 | 0.23 |
| 5s/30m | 147 | 1.44 | 0.15 |
| 5s/15m | 75 | 1.25 | 0.13 |
| 5s/5m | 172 | 0.87 | 0.07 |
| 5s/3m | 215 | 1.56 | 0.15 |
| 5s/m | 92 | 1.88 | 0.32 |
| 5s/30s | 67 | 2.09 | 0.57 |
| 5s/15s | 35 | 3.84 | 1.17 |
| SuppFig1-G | | | |
| GroupID | n | Total rolling time (s) | SEM |
| Naïve | 407 | 3.53 | 0.17 |
| 5s/30m | 147 | 4.79 | 0.33 |
| 5s/15m | 75 | 6.47 | 0.58 |
| 5s/5m | 172 | 11.00 | 0.34 |
| 5s/3m | 215 | 7.92 | 0.26 |
| 5s/m | 92 | 2.96 | 0.21 |
| 5s/30s | 67 | 1.72 | 0.13 |
| 5s/15s | 35 | 2.59 | 0.33 |
| SuppFig2-C | | | |
| GroupID | n | AUC | SEM |
| Naïve | 8 | 23.03 | 1.24 |
| Stim | 7 | 25.71 | 1.32 |
| SuppFig2-D | | | |
| GroupID | n | Critical Radius | SEM |
| Naïve | 8 | 50.13 | 3.41 |
| Stim | 7 | 52.29 | 4.09 |
| SuppFig2-E | | | |
| GroupID | n | Peak Intersection | SEM |
| Naïve | 8 | 47.75 | 2.84 |
| Stim | 7 | 52.57 | 3.63 |
| SuppFig3-A | | | |
| GroupID | n | Rolling probability (%) | 95% CI |

| | | | |
|---|---|---|---|
| No RNAi / No Dev. Stim. | 100 | 48.00 | 9.79 |
| No RNAi / Dev. Stim. | 100 | 85.00 | 7.00 |
| Dop1R1-RNAi | 100 | 70.00 | 8.98 |
| Dop1R2-RNAi (1) | 100 | 80.00 | 7.84 |
| Dop1R2-RNAi (2) | 100 | 62.00 | 9.51 |
| Dop2R-RNAi | 93 | 62.37 | 9.85 |
| dTk99R-RNAi | 100 | 74.00 | 8.60 |
| GABA-B-R1-RNAi | 100 | 74.00 | 8.60 |
| GABA-B-R2-RNAi | 100 | 86.00 | 6.80 |
| OAMB-RNAi (1) | 100 | 37.00 | 9.46 |
| OAMB-RNAi (2) | 100 | 13.00 | 6.59 |
| Octa2R-RNAi | 100 | 69.00 | 9.06 |
| OctB1R-RNAi | 100 | 96.00 | 3.84 |
| OctB2R-RNAi | 100 | 65.00 | 9.35 |
| OctB3R-RNAi | 100 | 51.00 | 9.80 |
| RYaR-RNAi | 70 | 100.00 | 0.00 |
| sNPFR-RNAi | 100 | 98.00 | 2.74 |
| TyrRI-RNAi | 100 | 70.00 | 8.98 |
| TyrRII-RNAi | 100 | 86.00 | 6.80 |
| SuppFig3-B | | | |
| GroupID | n | Latency (s) | SEM |
| No RNAi / No Dev. Stim. | 48 | 2.80 | 0.63 |
| No RNAi / Dev. Stim. | 85 | 0.94 | 0.07 |
| Dop1R1-RNAi | 70 | 1.26 | 0.12 |
| Dop1R2-RNAi (1) | 80 | 1.63 | 0.21 |
| Dop1R2-RNAi (2) | 62 | 1.13 | 0.36 |
| Dop2R-RNAi | 58 | 1.56 | 0.32 |
| dTk99R-RNAi | 74 | 0.66 | 0.05 |
| GABA-B-R1-RNAi | 74 | 1.08 | 0.12 |
| GABA-B-R2-RNAi | 86 | 0.84 | 0.06 |
| OAMB-RNAi (1) | 37 | 3.37 | 0.38 |
| OAMB-RNAi (2) | 13 | 2.44 | 0.79 |
| Octa2R-RNAi | 69 | 1.29 | 0.18 |
| OctB1R-RNAi | 96 | 1.40 | 0.11 |
| OctB2R-RNAi | 65 | 1.50 | 0.16 |
| OctB3R-RNAi | 51 | 1.54 | 0.57 |
| RYaR-RNAi | 70 | 0.81 | 0.07 |
| sNPFR-RNAi | 98 | 0.85 | 0.07 |

| | | | |
|---|---|---|---|
| TyrRI-RNAi | 70 | 1.94 | 0.47 |
| TyrRII-RNAi | 86 | 1.32 | 0.19 |
| SuppFig3-C | | | |
| GroupID | n | Total rolling time (s) | SEM |
| No RNAi / No Dev. Stim. | 48 | 2.38 | 0.56 |
| No RNAi / Dev. Stim. | 85 | 5.54 | 0.41 |
| Dop1R1-RNAi | 70 | 2.00 | 0.17 |
| Dop1R2-RNAi (1) | 80 | 3.07 | 0.23 |
| Dop1R2-RNAi (2) | 62 | 2.70 | 0.15 |
| Dop2R-RNAi | 58 | 2.44 | 0.28 |
| dTk99R-RNAi | 74 | 6.81 | 0.41 |
| GABA-B-R1-RNAi | 74 | 4.44 | 0.44 |
| GABA-B-R2-RNAi | 86 | 7.58 | 0.31 |
| OAMB-RNAi (1) | 37 | 1.21 | 0.16 |
| OAMB-RNAi (2) | 13 | 0.80 | 0.09 |
| Octa2R-RNAi | 69 | 2.87 | 0.21 |
| OctB1R-RNAi | 96 | 3.62 | 0.39 |
| OctB2R-RNAi | 65 | 2.28 | 0.27 |
| OctB3R-RNAi | 51 | 2.69 | 0.31 |
| RYaR-RNAi | 70 | 6.87 | 0.38 |
| sNPFR-RNAi | 98 | 6.50 | 0.34 |
| TyrRI-RNAi | 70 | 4.65 | 0.56 |
| TyrRII-RNAi | 86 | 4.67 | 0.42 |
| SuppFig4-A | | | |
| GroupID | n | Latency (s) | SEM |
| tbh-RNAi_1>- / Dev. Stim. | 292 | 5.23 | 0.26 |
| tbh-RNAi_1>- / No Dev. Stim. | 44 | 6.87 | 0.91 |
| tbh-RNAi_1>tdc2 / Dev. Stim. | 226 | 9.30 | 0.35 |
| tbh-RNAi_1>tdc2 / No Dev. Stim. | 83 | 8.48 | 0.58 |
| tbh-RNAi_2>- / Dev. Stim. | 163 | 0.84 | 0.18 |
| tbh-RNAi_2>- / No Dev. Stim. | 55 | 3.01 | 0.47 |
| tbh-RNAi_2>tdc2 / Dev. Stim. | 167 | 3.29 | 0.27 |
| tbh-RNAi_2>tdc2 / No Dev. Stim. | 70 | 3.97 | 0.37 |
| SuppFig4-B | | | |
| GroupID | n | Total rolling time (s) | SEM |
| tbh-RNAi_1>- / Dev. Stim. | 292 | 8.52 | 0.42 |
| tbh-RNAi_1>- / No Dev. Stim. | 44 | 1.16 | 0.14 |

| | | | |
|---|---|---|---|
| tbh-RNAi_1>tdc2 / Dev. Stim. | 226 | 5.16 | 0.31 |
| tbh-RNAi_1>tdc2 / No Dev. Stim. | 83 | 1.63 | 0.27 |
| tbh-RNAi_2>- / Dev. Stim. | 163 | 16.43 | 0.55 |
| tbh-RNAi_2>- / No Dev. Stim. | 55 | 5.01 | 0.52 |
| tbh-RNAi_2>tdc2 / Dev. Stim. | 167 | 11.36 | 0.56 |
| tbh-RNAi_2>tdc2 / No Dev. Stim. | 70 | 4.27 | 0.36 |
| SuppFig4-C | | | |
| GroupID | n | Latency (s) | SEM |
| attp2 / Dev. Stim. | 263 | 1.68 | 0.15 |
| attp2 / No Dev. Stim. | 144 | 4.24 | 0.45 |
| tdc2-GAL4 / Dev. Stim. | 265 | 1.25 | 0.19 |
| tdc2-GAL4 / No Dev. Stim. | 217 | 2.73 | 0.36 |
| SuppFig4-D | | | |
| GroupID | n | Total rolling time (s) | SEM |
| attp2 / Dev. Stim. | 263 | 14.28 | 0.45 |
| attp2 / No Dev. Stim. | 144 | 3.35 | 0.28 |
| tdc2-GAL4 / Dev. Stim. | 265 | 18.59 | 0.50 |
| tdc2-GAL4 / No Dev. Stim. | 217 | 5.78 | 0.34 |

Supplemental Table 4. Summary of statistical analysis per figure.

| Fig1-C Statistical Analysis | | | |
|---|---|---|---|
| Group1 | Group2 | Chi-Square | p value |
| Naïve | Stim. | 213.895 | 2E-48 |
| Fig1-D Statistical Analysis | | | |
| Group1 | Group2 | Welch's ANOVA (F-ratio) | p value |
| Naïve | Stim. | 54.819 | 1.19E-11 |
| Fig1-D Statistical Analysis | | | |
| Group1 | Group2 | Welch's ANOVA (F-ratio) | p value |
| Naïve | Stim. | 298.381 | 4.52E-48 |
| Fig1-F Statistical Analysis | | | |
| Group1 | Group2 | Chi-Square | p value |
| Naïve | 0-120h | 202.272 | 6.67E-46 |
| Naïve | 0-24h | 3.565 | 0.059 |
| Naïve | 24-48h | 2.437 | 0.1185 |
| Naïve | 48-72h | 10.373 | 0.0013 |
| Naïve | 72-96h | 53.992 | 2.01E-13 |
| Naïve | 96-120h | 75.205 | 4.24E-18 |
| 0-24h | 24-48h | 0.15 | 0.698 |
| 0-24h | 48-72h | 2.083 | 0.149 |
| 0-24h | 72-96h | 30.708 | 3E-08 |
| 0-24h | 96-120h | 45.286 | 1.7E-11 |
| 0-24h | 0-120h | 155.733 | 9.68E-36 |
| 24-48h | 48-72h | 3.413 | 0.0647 |
| 24-48h | 72-96h | 36.826 | 1.29E-09 |
| 24-48h | 96-120h | 53.997 | 2.01E-13 |
| 24-48h | 0-120h | 172.811 | 1.8E-39 |
| 48-72h | 72-96h | 14.93 | 0.000112 |
| 48-72h | 96-120h | 23.527 | 1.23E-06 |
| 48-72h | 0-120h | 111.897 | 3.76E-26 |
| 72-96h | 96-120h | 0.551 | 0.4577 |
| 72-96h | 0-120h | 48.837 | 2.78E-12 |
| 96-120h | 0-120h | 44.488 | 2.56E-11 |
| Fig1-G Statistical Analysis | | | |
| Group1 | Group2 | Chi-Square | p value |
| Naïve | 5s/30m | 0.909 | 0.3403 |
| Naïve | 5s/15m | 23.876 | 1.027E-06 |

| | | | |
|---|---|---|---|
| Naïve | 5s/5m | 191.485 | 1.51E-43 |
| Naïve | 5s/3m | 222.07 | 3.2E-50 |
| Naïve | 5s/m | 0.342 | 0.5589 |
| Naïve | 5s/30s | 13.199 | 0.0003 |
| Naïve | 5s/15s | 50.402 | 1.25E-12 |
| 5s/30m | 5s/15m | 15.236 | 0.00009489 |
| 5s/30m | 5s/5m | 140.492 | 2.08E-32 |
| 5s/30m | 5s/3m | 155.844 | 9.16E-36 |
| 5s/30m | 5s/m | 1.456 | 0.2276 |
| 5s/30m | 5s/30s | 14.536 | 0.0001374 |
| 5s/30m | 5s/15s | 46.17 | 6.51E-12 |
| 5s/15m | 5s/5m | 42.074 | 8.79E-11 |
| 5s/15m | 5s/3m | 41.942 | 9.4E-11 |
| 5s/15m | 5s/m | 21.219 | 4.097E-06 |
| 5s/15m | 5s/30s | 42.791 | 6.09E-11 |
| 5s/15m | 5s/15s | 78.472 | 8.12E-19 |
| 5s/5m | 5s/3m | 0.303 | 0.5818 |
| 5s/5m | 5s/m | 147.949 | 4.87E-34 |
| 5s/5m | 5s/30s | 199.816 | 2.29E-45 |
| 5s/5m | 5s/15s | 265.056 | 1.36E-59 |
| 5s/3m | 5s/m | 162.06 | 4.01E-37 |
| 5s/3m | 5s/30s | 220.168 | 8.31E-50 |
| 5s/3m | 5s/15s | 291.122 | 2.83E-65 |
| 5s/m | 5s/30s | 5.868 | 0.0152 |
| 5s/m | 5s/15s | 28.752 | 8.228E-08 |
| 5s/30s | 5s/15s | 9.249 | 0.002347 |
| Fig2-B Statistical Analysis | | | |
| Group1 | Group2 | Chi-Square | p-value |
| Omnibus Kruskall-Wallis Test (Ordinal Analysis) | - | 28.8255 | 7.82E-08 |
| Fig2-C Statistical Analysis | | | |
| Group1 | Group2 | Chi-Square | p-value |
| Omnibus Kruskall-Wallis Test (Ordinal Analysis) | - | 56.0921 | 6.92E-14 |
| Naïve-RollingProbability | Stim-RollingProbability | 27.614 | 1.48E-07 |
| Naïve-Latency Kruskall-Wallis Test | Stim-Latency | 30.3269 | 4E-08 |
| Fig2-D Statistical Analysis | | | |
| Group1 | Group2 | Chi-Square | p-value |

| | | | |
|---|---|---|---|
| Omnibus Nominal Logistic Regression | Whole Model | 389.7664 | 3.65E-84 |
| Omnibus Nominal Logistic Regression | Strength (mN) | 334.2676 | 1.13E-74 |
| Omnibus Nominal Logistic Regression | Group | 103.7476 | 2.3E-24 |
| Omnibus Nominal Logistic Regression | Strength (mN)*Group | 14.48805 | 0.000141 |
| Naïve-20mN | Stim.-20mN | 10.231 | 0.0014 |
| Naïve-40mN | Stim.-40mN | 4.465 | 0.0346 |
| Naïve-60mN | Stim.-60mN | 20.676 | 5.44E-06 |
| Naïve-80mN | Stim.-80mN | 13.115 | 0.000293 |
| Naïve-100mN | Stim.-100mN | 23.032 | 1.59E-06 |
| Naïve-120mN | Stim.-120mN | 19.972 | 7.86E-06 |
| Naïve-150mN | Stim.-150mN | 16.612 | 4.59E-05 |
| Fig2-E Statistical Analysis | | | |
| Group1 | Group2 | Chi-Square | p-value |
| Omnibus Nominal Logistic Regression | Whole Model | 382.8708 | 1.14E-82 |
| Omnibus Nominal Logistic Regression | Group | 58.24981 | 2.31E-14 |
| Omnibus Nominal Logistic Regression | HCl Concentration (%) | 353.2868 | 8.15E-79 |
| Omnibus Nominal Logistic Regression | Group*HCl Concentration (%) | 18.96636 | 1.33E-05 |
| Naïve-0.1% | Stim.-0.1% | 1.396 | 0.2373 |
| Naïve-0.5% | Stim.-0.5% | 38.861 | 4.55E-10 |
| Naïve-1% | Stim.-1% | 5.826 | 0.01579 |
| Naïve-2.5% | Stim.-2.5% | 4.672 | 0.03066 |
| Naïve-5% | Stim.-5% | 18.277 | 1.91E-05 |
| Naïve-7.5% | Stim.-7.5% | 11.787 | 0.000597 |
| Naïve-10% | Stim.-10% | 1.088 | 0.2969 |
| Fig2-F Statistical Analysis | | | |
| Group1 | Group2 | Chi-Square | p-value |
| Omnibus Nominal Logistic Regression | Whole Model | 316.4343 | 2.76E-68 |
| Omnibus Nominal Logistic Regression | Group | 30.17688 | 3.94E-08 |
| Omnibus Nominal Logistic Regression | Irradiance (uW/mm2) | 297.4503 | 1.18E-66 |
| Omnibus Nominal Logistic Regression | Group*Irradiance (uW/mm2) | 4.696662 | 0.03022 |

| | | | |
|---|---|---|---|
| Naive-0.01 | Stim.-0.01 | 0.925 | 0.3361 |
| Naive-0.03 | Stim.-0.03 | 8.276 | 0.004 |
| Naive-0.04 | Stim.-0.04 | 9.094 | 0.0026 |
| Naive-0.07 | Stim.-0.07 | 4 | 0.0455 |
| Naive-0.1 | Stim.-0.1 | 36.932 | 1.22E-09 |
| Fig2-I Statistical Analysis | | | |
| Group1 | Group2 | Mann-Whithney (Z approx.) | p-value |
| Naïve-0.1 | Stim.-0.1 | 3.198 | 0.0014 |
| Naïve-0.3 | Stim.-0.3 | 2.91 | 0.0036 |
| Naïve-0.5 | Stim.-0.5 | 3.0001 | 0.0027 |
| Naïve-0.9 | Stim.-0.9 | 2.985 | 0.0028 |
| Naïve-1.4 | Stim.-1.4 | 2.641 | 0.0075 |
| Group1 | Group2 | F-Ratio | p-value |
| Omnibus General Regression | Group | 51.41253 | 9.45E-11 |
| Omnibus General Regression | Irrandiance (uW/mm2) | 7.587566 | 0.00689 |
| Omnibus General Regression | Irrandiance (uW/mm2)*Group | 0.474737 | 0.492 |
| Fig3-A Statistical Analysis | | | |
| Group1 | Group2 | Chi Square | p-value |
| Omnibus ChiSquare Test | - | - | 5.56E-70 |
| OAMB-RNAi_1 / Dev. Stim. | No RNAi / Dev. Stim. | 237.996 | 1.08E-53 |
| OAMB-RNAi_1 / No Dev. Stim. | No RNAi / Dev. Stim. | 363.046 | 6.11E-81 |
| OAMB-RNAi_2 / Dev. Stim. | No RNAi / Dev. Stim. | 537.538 | 6.5E-119 |
| OAMB-RNAi_2 / No Dev. Stim. | No RNAi / Dev. Stim. | 263.6 | 2.82E-59 |
| No RNAi / No Dev. Stim. | No RNAi / Dev. Stim. | 335.938 | 4.89E-75 |
| OAMB-RNAi_1 / Dev. Stim. | No RNAi / No Dev. Stim. | 0.687 | 0.4072 |
| OAMB-RNAi_2 / No Dev. Stim. | No RNAi / No Dev. Stim. | 0.133 | 0.7155 |
| OAMB-RNAi_1 / No Dev. Stim. | No RNAi / No Dev. Stim. | 15.349 | 8.94E-05 |
| OAMB-RNAi_2 / Dev. Stim. | No RNAi / No Dev. Stim. | 63.924 | 1.29E-15 |
| OAMB-RNAi_1 / No Dev. Stim. | OAMB-RNAi_1 / Dev. Stim. | 16.749 | 4.27E-05 |
| OAMB-RNAi_2 / No Dev. Stim. | OAMB-RNAi_1 / Dev. Stim. | 1.037 | 0.3084 |
| OAMB-RNAi_2 / Dev. Stim. | OAMB-RNAi_1 / Dev. Stim. | 56.648 | 5.21E-14 |
| OAMB-RNAi_2 / No Dev. Stim. | OAMB-RNAi_1 / No Dev. Stim. | 9.291 | 0.002303 |

| | | | |
|---|---|---|---|
| OAMB-RNAi_2 / Dev. Stim. | OAMB-RNAi_1 / No Dev. Stim. | 9.529 | 0.002023 |
| OAMB-RNAi_2 / No Dev. Stim. | OAMB-RNAi_2 / Dev. Stim. | 40.533 | 1.93E-10 |
| Fig3-B Statistical Analysis | | | |
| Group1 | Group2 | Mann-Whithney (Z approx.) | p-value |
| OAMB-RNAi_1 / Dev. Stim. | No RNAi / Dev. Stim. | 10.86452 | 1.70E-27 |
| OAMB-RNAi_2 / No Dev. Stim. | No RNAi / Dev. Stim. | 10.62599 | 2.26E-26 |
| OAMB-RNAi_2 / Dev. Stim. | No RNAi / Dev. Stim. | 8.33072 | 8.03E-17 |
| OAMB-RNAi_1 / No Dev. Stim. | No RNAi / Dev. Stim. | 8.02127 | 1.05E-15 |
| No RNAi / No Dev. Stim. | No RNAi / Dev. Stim. | 7.44065 | 1.00E-13 |
| OAMB-RNAi_2 / No Dev. Stim. | No RNAi / No Dev. Stim. | 4.54948 | 5.38E-06 |
| OAMB-RNAi_2 / Dev. Stim. | No RNAi / No Dev. Stim. | 3.25508 | 1.13E-03 |
| OAMB-RNAi_1 / Dev. Stim. | No RNAi / No Dev. Stim. | 3.4262 | 6.12E-04 |
| OAMB-RNAi_1 / No Dev. Stim. | No RNAi / No Dev. Stim. | 3.00185 | 2.68E-03 |
| OAMB-RNAi_2 / No Dev. Stim. | OAMB-RNAi_1 / Dev. Stim. | 2.11979 | 3.40E-02 |
| OAMB-RNAi_2 / Dev. Stim. | OAMB-RNAi_1 / Dev. Stim. | 0.84857 | 3.96E-01 |
| OAMB-RNAi_2 / No Dev. Stim. | OAMB-RNAi_1 / No Dev. Stim. | 0.6884 | 4.91E-01 |
| OAMB-RNAi_2 / No Dev. Stim. | OAMB-RNAi_2 / Dev. Stim. | 0.66353 | 5.07E-01 |
| OAMB-RNAi_1 / No Dev. Stim. | OAMB-RNAi_1 / Dev. Stim. | 0.52307 | 6.01E-01 |
| OAMB-RNAi_2 / Dev. Stim. | OAMB-RNAi_1 / No Dev. Stim. | 0.29702 | 7.66E-01 |
| Fig3-C Statistical Analysis | | | |
| Group1 | Group2 | Mann-Whithney (Z approx.) | p-value |
| OAMB-RNAi_1 / Dev. Stim. | No RNAi / No Dev. Stim. | 2.8348 | 4.58E-03 |
| OAMB-RNAi_2 / No Dev. Stim. | OAMB-RNAi_2 / Dev. Stim. | 1.9632 | 4.96E-02 |
| OAMB-RNAi_2 / No Dev. Stim. | OAMB-RNAi_1 / No Dev. Stim. | 1.0307 | 3.03E-01 |
| OAMB-RNAi_2 / Dev. Stim. | OAMB-RNAi_1 / No Dev. Stim. | -0.2971 | 7.66E-01 |
| OAMB-RNAi_2 / No Dev. Stim. | No RNAi / No Dev. Stim. | -1.0609 | 2.89E-01 |

| | | | |
|---|---|---|---|
| OAMB-RNAi_1 / No Dev. Stim. | No RNAi / No Dev. Stim. | -2.3802 | 1.73E-02 |
| OAMB-RNAi_1 / No Dev. Stim. | OAMB-RNAi_1 / Dev. Stim. | -3.5503 | 3.85E-04 |
| OAMB-RNAi_2 / No Dev. Stim. | OAMB-RNAi_1 / Dev. Stim. | -3.4899 | 4.83E-04 |
| OAMB-RNAi_2 / Dev. Stim. | OAMB-RNAi_1 / Dev. Stim. | -4.544 | 5.52E-06 |
| OAMB-RNAi_2 / Dev. Stim. | No RNAi / No Dev. Stim. | -3.2005 | 1.37E-03 |
| OAMB-RNAi_1 / Dev. Stim. | No RNAi / Dev. Stim. | -10.0359 | 1.06E-23 |
| OAMB-RNAi_1 / No Dev. Stim. | No RNAi / Dev. Stim. | -10.0373 | 1.04E-23 |
| OAMB-RNAi_2 / Dev. Stim. | No RNAi / Dev. Stim. | -10.1032 | 5.34E-24 |
| OAMB-RNAi_2 / No Dev. Stim. | No RNAi / Dev. Stim. | -11.8011 | 3.85E-32 |
| No RNAi / No Dev. Stim. | No RNAi / Dev. Stim. | -13.7203 | 7.68E-43 |
| Fig3-F Statistical Analysis | | | |
| Group1 | Group2 | Mann-Whitney (Z approx.) | p-value |
| No RNAi / Stim. | No RNAi / Naïve | 2.73526 | 0.0062 |
| OAMB-RNAi / Naïve | No RNAi / Naïve | -0.11339 | 0.9097 |
| OAMB-RNAi / Stim. | No RNAi / Naïve | -0.61237 | 0.5403 |
| OAMB-RNAi / Stim. | OAMB-RNAi / Naïve | -0.69402 | 0.4877 |
| OAMB-RNAi / Naïve | No RNAi / Stim. | -2.81691 | 0.0048 |
| OAMB-RNAi / Stim. | No RNAi / Stim. | -3.35548 | 0.0008 |
| Fig3-H Statistical Analysis | | | |
| Group1 | Group2 | ANOVA (F-ratio) | p-value |
| ddaC / Naïve | ddaC / Stim. | 45.098 | 2.15E-05 |
| v'ada / Naïve | v'ada / Stim. | 0.5113 | 0.4883 |
| vdaB / Naïve | vdaB / Stim. | 0.8265 | 0.3812 |
| Fig4-A Statistical Analysis | | | |
| Group1 | Group2 | Chi-Square | p value |
| tbh-RNAi_2>- / Naïve | tbh-RNAi_2>tdc2 / Naïve | 1.912 | 0.1668 |
| tbh-RNAi_2>- / Naïve | tbh-RNAi_2>- / Stim. | 100.033 | 1.5E-23 |
| tbh-RNAi_2>- / Naïve | tbh-RNAi_2>tdc2 / Stim. | 41.2 | 1.37E-10 |
| tbh-RNAi_2>tdc2 / Naïve | tbh-RNAi_2>- / Stim. | 154.996 | 1.4E-35 |
| tbh-RNAi_2>tdc2 / Naïve | tbh-RNAi_2>tdc2 / Stim. | 77.201 | 1.54E-18 |
| tbh-RNAi_2>- / Stim. | tbh-RNAi_2>tdc2 / Stim. | 20.518 | 5.909E-06 |

| | | | |
|---|---|---|---|
| Tbh-RNAi_1>- / Naïve | Tbh-RNAi_1>tdc2/ Naïve | 0.24 | 0.6241 |
| Tbh-RNAi_1>- / Naïve | Tbh-RNAi_1>- / Stim. | 371.164 | 1.04E-82 |
| Tbh-RNAi_1>- / Naïve | Tbh-RNAi_1>tdc2/ Stim. | 210.05 | 1.34E-47 |
| Tbh-RNAi_1>tdc2/ Naïve | Tbh-RNAi_1>- / Stim. | 460.359 | 4E-102 |
| Tbh-RNAi_1>tdc2/ Naïve | Tbh-RNAi_1>tdc2/ Stim. | 259.839 | 1.86E-58 |
| Tbh-RNAi_1>- / Stim. | Tbh-RNAi_1>tdc2/ Stim. | 27.816 | 1.34E-07 |
| Tbh-RNAi_1>- / Stim. | tbh-RNAi_2>- / Stim. | 2.533 | 0.11147 |
| Tbh-RNAi_1>- / Naïve | tbh-RNAi_2>- / Naïve | 18.993 | 1.31E-05 |
| Tbh-RNAi_1>tdc2 / Stim. | tbh-RNAi_2>tdc2 / Stim. | 3.531 | 0.060231 |
| Tbh-RNAi_1>tdc2 / Naïve | tbh-RNAi_2>tdc2/ Naïve | 10.766 | 0.001034 |
| Tbh-RNAi_1>tdc2 / Stim. | tbh-RNAi_2>- / Stim. | 9.001 | 0.0027 |
| Fig4-C Statistical Analysis | | | |
| Group1 | Group2 | Chi-Square | p value |
| attp2 / No Dev. Stim. | attp2 / Dev. Stim. | 183.511 | 8.30E-42 |
| tdc2-GAL4 / No Dev. Stim. | tdc2-GAL4 / Dev. Stim. | 16.626 | 4.55E-05 |
| attp2 / No Dev. Stim. | tdc2-GAL4 / No Dev. Stim. | 5.866 | 0.154 |
| attp2 / Dev. Stim. | tdc2-GAL4 / Dev. Stim. | 70.8 | 3.95E-17 |
| attp2 / No Dev. Stim. | tdc2-GAL4 / Dev. Stim. | 38.948 | 4.35E-10 |
| attp2 / Dev. Stim. | tdc2-GAL4 / No Dev. Stim. | 142.411 | 4.78E-33 |
| Fig5-B Statistical Analysis | | | |
| Group1 | Group2 | Chi-Square | p value |
| 0mM | 5mM | 17.372 | 3.07E-05 |
| 0mM | 25mM | 23.134 | 1.51E-06 |
| 0mM | 50mM | 31.442 | 2.06E-08 |
| 5mM | 25mM | 0.502 | 4.79E-01 |
| 5mM | 50mM | 3.051 | 0.0807 |
| 25mM | 50mM | 1.137 | 0.2863 |
| Fig5-C Statistical Analysis | | | |
| Group1 | Group2 | Chi-Square | p-value |
| Omnibus Kruskall-Wallis Test (Ordinal Analysis) | - | 9.1743 | 0.0025 |
| Fig5-D Statistical Analysis | | | |
| Group1 | Group2 | Chi-Square | p-value |
| Omnibus Kruskall-Wallis Test (Ordinal Analysis) | - | 9.8532 | 0.0017 |
| Naïve-RollingProbability | Stim-RollingProbability | 3.7895 | 0.05157 |

| | | | |
|---|---|---|---|
| Naïve-Latency Kruskall-Wallis Test | Stim-Latency | 6.7164 | 0.0096 |
| Group1 | Group2 | Mann-Whithney (Z approx.) | p-value |
| Untreated-0.1 | 5mM OA-0.1 | 2.00321 | 0.0226 |
| Untreated-0.5 | 5mM OA-0.5 | 2.14359 | 0.016 |
| Untreated-1.4 | 5mM OA-1.4 | 1.70084 | 0.0445 |
| Group1 | Group2 | F-Ratio | p-value |
| Omnibus General Regression | Condition | 13.69553 | 0.0005 |
| Omnibus General Regression | LED | 6.944522 | 0.0108 |
| Omnibus General Regression | LED*Condition | 0.906575 | 0.345 |
| Fig6-C Statistical Analysis | | | |
| Group1 | Group2 | Mann-Whitney (Z approx.) | p-value |
| VUM-Stim | attp2-Naïve | 3.01664 | 0.0026 |
| tdc2-tsh-FLP-Stim. | attp2-Naïve | 2.9841 | 0.0028 |
| tdc2-Stim. | attp2-Naïve | 2.86038 | 0.0042 |
| VUM-Stim | attp2-Stim. | 2.75145 | 0.0059 |
| tdc2-tsh-FLP-Stim. | attp2-Stim. | 2.72648 | 0.0064 |
| tdc2-Stim. | attp2-Stim. | 2.59534 | 0.0094 |
| VUM-Stim | VUM-Naïve | 3.01963 | 0.0025 |
| VUM-Stim | tdc2-Naïve | 3.00536 | 0.0027 |
| tdc2-tsh-FLP-Stim. | tdc2-Naïve | 2.98676 | 0.0028 |
| tdc2-Stim. | tdc2-Naïve | 2.87305 | 0.0041 |
| tdc2-tsh-FLP-Stim. | tdc2-tsh-FLP-Naïve | 2.73224 | 0.0063 |
| VUM-Stim | tdc2-tsh-FLP-Naïve | 2.71723 | 0.0066 |
| tdc2-tsh-FLP-Stim. | tdc2-tsh-gal80-Stim. | 2.02874 | 0.0425 |
| VUM-Stim | tdc2-tsh-gal80-Stim. | 1.99945 | 0.0456 |
| tdc2-Stim. | tdc2-tsh-gal80-Stim. | 1.88756 | 0.0591 |
| tdc2-tsh-FLP-Stim. | tdc2-tsh-gal80-Naïve | 1.75249 | 0.0797 |
| VUM-Stim | tdc2-tsh-gal80-Naïve | 1.71559 | 0.0862 |
| tdc2-Stim. | tdc2-tsh-gal80-Naïve | 1.60597 | 0.1083 |
| tdc2-tsh-gal80-Naïve | attp2-Naïve | 1.07068 | 0.2843 |
| tdc2-tsh-gal80-Naïve | attp2-Stim. | 0.7914 | 0.4287 |
| tdc2-tsh-gal80-Stim. | attp2-Naïve | 0.74327 | 0.4573 |
| tdc2-tsh-gal80-Stim. | attp2-Stim. | 0.46097 | 0.6448 |
| attp2-Stim. | attp2-Naïve | 0.37963 | 0.7042 |
| tdc2-tsh-FLP-Naïve | tdc2-Naïve | 0.30449 | 0.7608 |

| | | | |
|---|---|---|---|
| tdc2-tsh-FLP-Stim. | tdc2-Stim. | 0.17231 | 0.8632 |
| VUM-Stim | tdc2-Stim. | 0.09182 | 0.9268 |
| tdc2-tsh-FLP-Naïve | attp2-Naïve | -0.04891 | 0.961 |
| VUM-Stim | tdc2-tsh-FLP-Stim. | -0.08416 | 0.9329 |
| VUM-Naïve | tdc2-Naïve | -0.08702 | 0.9307 |
| tdc2-tsh-gal80-Stim. | tdc2-tsh-gal80-Naïve | -0.27781 | 0.7812 |
| VUM-Naïve | tdc2-tsh-FLP-Naïve | -0.39622 | 0.6919 |
| tdc2-tsh-FLP-Naïve | attp2-Stim. | -0.32965 | 0.7417 |
| tdc2-Naïve | attp2-Naïve | -0.38722 | 0.6986 |
| VUM-Naïve | attp2-Naïve | -0.46951 | 0.6387 |
| tdc2-tsh-FLP-Naïve | tdc2-tsh-gal80-Stim. | -0.72933 | 0.4658 |
| tdc2-Naïve | attp2-Stim. | -0.68897 | 0.4908 |
| VUM-Naïve | attp2-Stim. | -0.75818 | 0.4483 |
| tdc2-tsh-FLP-Naïve | tdc2-tsh-gal80-Naïve | -1.00255 | 0.3161 |
| tdc2-Naïve | tdc2-tsh-gal80-Stim. | -1.01254 | 0.3113 |
| VUM-Naïve | tdc2-tsh-gal80-Stim. | -1.08509 | 0.2779 |
| tdc2-Naïve | tdc2-tsh-gal80-Naïve | -1.29203 | 0.1963 |
| VUM-Naïve | tdc2-tsh-gal80-Naïve | -1.35442 | 0.1756 |
| tdc2-tsh-FLP-Naïve | tdc2-Stim. | -2.60052 | 0.0093 |
| VUM-Naïve | tdc2-Stim. | -2.89202 | 0.0038 |
| VUM-Naïve | tdc2-tsh-FLP-Stim. | -3.00531 | 0.0027 |
| Fig6-D Statistical Analysis | | | |
| Group1 | Group2 | Mann-Whitney (Z approx.) | p-value |
| VUM-Stim | tdc2-tsh-gal80-Stim. | -5.2245 | 1.75E-07 |
| tdc2-tsh-FLP-Stim. | tdc2-tsh-gal80-Stim. | -5.17703 | 2.26E-07 |
| tdc2-tsh-FLP-Stim. | tdc2-tsh-gal80-Naïve | -4.97997 | 6.36E-07 |
| VUM-Stim | tdc2-tsh-gal80-Naïve | -4.95638 | 7.18E-07 |
| VUM-Stim | attp2-Naïve | -4.91647 | 8.81E-07 |
| tdc2-tsh-FLP-Stim. | attp2-Naïve | -4.8616 | 1.16E-06 |
| VUM-Stim | tdc2-Naïve | -4.71701 | 2.39E-06 |
| tdc2-tsh-FLP-Stim. | tdc2-Naïve | -4.70075 | 2.59E-06 |
| VUM-Stim | tdc2-tsh-FLP-Naïve | -4.43202 | 9.34E-06 |
| VUM-Stim | attp2-Stim. | -4.28965 | 1.79E-05 |
| tdc2-tsh-FLP-Stim. | tdc2-tsh-FLP-Naïve | -4.26816 | 1.97E-05 |
| tdc2-Stim. | tdc2-tsh-gal80-Stim. | -4.24384 | 2.20E-05 |
| tdc2-Stim. | attp2-Naïve | -4.06919 | 4.72E-05 |

| | | | |
|---|---|---|---|
| tdc2-tsh-FLP-Stim. | attp2-Stim. | -4.03964 | 5.35E-05 |
| tdc2-Stim. | tdc2-tsh-gal80-Naïve | -4.02939 | 5.59E-05 |
| tdc2-Stim. | tdc2-Naïve | -3.8109 | 1.39E-04 |
| VUM-Stim | VUM-Naïve | -3.64891 | 2.63E-04 |
| VUM-Naïve | tdc2-tsh-FLP-Stim. | 3.43525 | 5.92E-04 |
| tdc2-tsh-FLP-Naïve | tdc2-Stim. | 3.36958 | 7.53E-04 |
| tdc2-Stim. | attp2-Stim. | -3.23397 | 1.22E-03 |
| VUM-Naïve | tdc2-Stim. | 2.62012 | 8.79E-03 |
| VUM-Naïve | attp2-Naïve | -1.69619 | 8.99E-02 |
| VUM-Naïve | tdc2-tsh-gal80-Stim. | -1.44785 | 1.48E-01 |
| VUM-Naïve | tdc2-tsh-gal80-Naïve | -1.34853 | 1.78E-01 |
| VUM-Stim | tdc2-Stim. | -1.30365 | 1.92E-01 |
| VUM-Naïve | tdc2-Naïve | -1.22109 | 2.22E-01 |
| tdc2-tsh-FLP-Naïve | attp2-Naïve | -1.17982 | 2.38E-01 |
| attp2-Stim. | attp2-Naïve | -1.05038 | 2.94E-01 |
| tdc2-tsh-FLP-Naïve | tdc2-tsh-gal80-Stim. | -0.88804 | 3.75E-01 |
| tdc2-tsh-FLP-Naïve | tdc2-tsh-gal80-Naïve | -0.85228 | 3.94E-01 |
| tdc2-tsh-gal80-Stim. | attp2-Stim. | 0.82219 | 4.11E-01 |
| tdc2-tsh-FLP-Stim. | tdc2-Stim. | -0.82162 | 4.11E-01 |
| tdc2-tsh-FLP-Naïve | tdc2-Naïve | -0.72361 | 4.69E-01 |
| tdc2-tsh-gal80-Naïve | attp2-Stim. | 0.7042 | 4.81E-01 |
| VUM-Naïve | tdc2-tsh-FLP-Naïve | -0.64516 | 5.19E-01 |
| VUM-Naïve | attp2-Stim. | -0.62049 | 5.35E-01 |
| tdc2-Naïve | attp2-Stim. | 0.58564 | 5.58E-01 |
| VUM-Stim | tdc2-tsh-FLP-Stim. | -0.58176 | 5.61E-01 |
| tdc2-Naïve | attp2-Naïve | -0.54238 | 5.88E-01 |
| tdc2-tsh-gal80-Naïve | attp2-Naïve | -0.49086 | 6.24E-01 |
| tdc2-tsh-gal80-Stim. | attp2-Naïve | -0.40326 | 6.87E-01 |
| tdc2-Naïve | tdc2-tsh-gal80-Stim. | -0.1435 | 8.86E-01 |
| tdc2-tsh-gal80-Stim. | tdc2-tsh-gal80-Naïve | 0.06811 | 9.46E-01 |
| tdc2-Naïve | tdc2-tsh-gal80-Naïve | -0.05686 | 9.55E-01 |
| tdc2-tsh-FLP-Naïve | attp2-Stim. | 0.04555 | 9.64E-01 |
| Fig6-H Statistical Analysis | | | |
| Group1 | Group2 | Mann-Whitney (Z approx.) | p-value |
| Naïve | Stim. | 3.752 | 0.0002 |
| Fig6-H Statistical Analysis | | | |

| Group1 | Group2 | One-Way ANOVA (F Ratio) | p-value |
|---|---|---|---|
| Naïve | Stim. | 7.5945 | 0.009 |
| SuppFig1-A Statistical Analysis | | | |
| Group1 | Group2 | Mann-Whitney (Z approx.) | p value |
| Naïve | 0-120h | 9.38718 | 6.16E-21 |
| 24-48h | 0-120h | 8.73828 | 2.37E-18 |
| 96-120h | 0-120h | 7.15464 | 8.39E-13 |
| 48-72h | 0-120h | 7.06122 | 1.65E-12 |
| Naïve | 72-96h | 5.72425 | 1.04E-08 |
| 72-96h | 0-120h | 4.43137 | 9.36E-06 |
| Naïve | 96-120h | 4.637 | 3.54E-06 |
| Naïve | 48-72h | 3.55023 | 0.000385 |
| 96-120h | 72-96h | 2.20834 | 0.02722 |
| Naïve | 0-24h | 2.67243 | 0.00753 |
| Naïve | 24-48h | 2.30678 | 0.02107 |
| 24-48h | 0-24h | 0.677 | 0.4984 |
| 48-72h | 0-24h | -0.82875 | 0.4072 |
| 96-120h | 48-72h | -0.85066 | 0.395 |
| 48-72h | 24-48h | -1.76619 | 0.07736 |
| 96-120h | 0-24h | -1.78457 | 0.07433 |
| 72-96h | 48-72h | -2.88399 | 0.003927 |
| 96-120h | 24-48h | -2.54658 | 0.01088 |
| 72-96h | 0-24h | -3.54258 | 0.000396 |
| 72-96h | 24-48h | -4.4814 | 7.42E-06 |
| 0-120h | 0-24h | -7.69945 | 1.37E-14 |
| SuppFig1-B Statistical Analysis | | | |
| Group1 | Group2 | Mann-Whitney (Z approx.) | p value |
| 0-120h | 0-24h | 12.6822 | 7.42E-37 |
| 96-120h | 24-48h | 8.7273 | 2.61E-18 |
| 96-120h | 48-72h | 7.3248 | 2.39E-13 |
| 96-120h | 0-24h | 6.1681 | 6.91E-10 |
| 72-96h | 24-48h | 5.1989 | 2.01E-07 |
| 96-120h | 72-96h | 4.2825 | 1.85E-05 |

| | | | |
|---|---|---|---|
| 72-96h | 48-72h | 3.8989 | 9.66E-05 |
| 72-96h | 0-24h | 2.6392 | 0.008309 |
| 48-72h | 24-48h | 0.9716 | 0.3312 |
| Naïve | 24-48h | -0.5133 | 0.6077 |
| 48-72h | 0-24h | -1.1435 | 0.2528 |
| Naïve | 48-72h | -1.3092 | 0.1905 |
| 24-48h | 0-24h | -2.1704 | 0.02998 |
| Naïve | 0-24h | -2.3641 | 0.01807 |
| Naïve | 72-96h | -5.134 | 2.84E-07 |
| Naïve | 96-120h | -8.076 | 6.69E-16 |
| 96-120h | 0-120h | -8.8808 | 6.64E-19 |
| 72-96h | 0-120h | -12.2472 | 1.74E-34 |
| 48-72h | 0-120h | -13.2554 | 4.2E-40 |
| Naïve | 0-120h | -13.3048 | 2.17E-40 |
| 24-48h | 0-120h | -14.8065 | 1.33E-49 |
| SuppFig1-C Statistical Analysis | | | |
| Group1 | Group2 | Chi-Square | p value |
| Omnibus Chi-Square Test | - | 466.792 | 1.17E-98 |
| Naïve | 0.14 | 6.221 | 0.0126 |
| Naïve | 0.35 | 166.913 | 3.5E-38 |
| Naïve | 1.41 | 184.115 | 6.12E-42 |
| Naïve | 2.82 | 122.17 | 2.12E-28 |
| Naïve | 5.64 | 185.962 | 2.42E-42 |
| 0.14 | 0.35 | 92.23 | 7.72E-22 |
| 0.14 | 1.41 | 113.093 | 2.06E-26 |
| 0.14 | 2.82 | 71.772 | 2.42E-17 |
| 0.14 | 5.64 | 100.254 | 1.34E-23 |
| 0.35 | 1.41 | 4.388 | 0.03619 |
| 0.35 | 2.82 | 0.024 | 0.8774 |
| 0.35 | 5.64 | 0.004 | 0.951 |
| 1.41 | 2.82 | 3.912 | 0.0479 |
| 1.41 | 5.64 | 4.559 | 0.0327 |
| 2.82 | 5.64 | 0.045 | 0.8313 |
| SuppFig1-D Statistical Analysis | | | |
| Group1 | Group2 | Mann-Whitney (Z approx.) | p value |
| '1.41' | '0' | -12.3958 | 2.75E-35 |
| '5.64' | '0' | -11.3645 | 6.28E-30 |

| | | | |
|---|---|---|---|
| '1.41' | '0.14' | -10.1748 | 2.57E-24 |
| '0' | '0.35' | 9.0892 | 9.97E-20 |
| '2.82' | '0' | -9.034 | 1.65E-19 |
| '5.64' | '0.14' | -8.9368 | 4.01E-19 |
| '2.82' | '0.14' | -7.4919 | 6.79E-14 |
| '0.35' | '0.14' | -7.1997 | 6.04E-13 |
| '1.41' | '0.35' | -6.0063 | 1.9E-09 |
| '2.82' | '1.41' | 3.9169 | 8.97E-05 |
| '5.64' | '1.41' | 2.9198 | 0.003503 |
| '5.64' | '0.35' | -2.7025 | 0.006882 |
| '2.82' | '0.35' | -1.4936 | 0.1353 |
| '5.64' | '2.82' | -0.9876 | 0.3233 |
| '0' | '0.14' | 0.6207 | 0.5348 |
| SuppFig1-E Statistical Analysis | | | |
| Group1 | Group2 | Mann-Whitney (Z approx.) | p value |
| '5.64' | '0' | 15.7346 | 8.76E-56 |
| '1.41' | '0' | 14.9244 | 2.29E-50 |
| '2.82' | '0' | 13.6032 | 3.83E-42 |
| '0' | '0.35' | -13.0822 | 4.16E-39 |
| '1.41' | '0.14' | 11.3991 | 4.23E-30 |
| '5.64' | '0.14' | 10.5551 | 4.81E-26 |
| '2.82' | '0.14' | 10.292 | 7.66E-25 |
| '0.35' | '0.14' | 7.5806 | 3.44E-14 |
| '1.41' | '0.35' | 6.9491 | 3.68E-12 |
| '2.82' | '0.35' | 6.1592 | 7.31E-10 |
| '0' | '0.14' | -5.0465 | 4.5E-07 |
| '5.64' | '0.35' | 4.244 | 2.2E-05 |
| '5.64' | '1.41' | -3.1742 | 0.001503 |
| '5.64' | '2.82' | -3.1523 | 0.00162 |
| '2.82' | '1.41' | 0.8162 | 0.4144 |
| SuppFig1-F Statistical Analysis | | | |
| Group1 | Group2 | Mann-Whitney (Z approx.) | p value |
| Naïve | 5s/5m | 8.29104 | 1.12E-16 |
| 5s/5m | 5s/1m | -6.72359 | 1.77E-11 |
| 5s/5m | 5s/3m | -5.23628 | 1.64E-07 |

| | | | |
|---|---|---|---|
| Naïve | 5s/30m | 4.33539 | 1.46E-05 |
| 5s/15s | 5s/5m | 4.32269 | 1.54E-05 |
| 5s/15m | 5s/5m | 3.93135 | 8.45E-05 |
| 5s/30m | 5s/1m | -3.46713 | 0.000526 |
| 5s/30s | 5s/5m | 3.24385 | 0.001179 |
| Naïve | 5s/3m | 3.23036 | 0.001236 |
| 5s/30m | 5s/5m | 3.15901 | 0.001583 |
| 5s/30s | 5s/1m | -2.70011 | 0.006932 |
| Naïve | 5s/30s | 2.61238 | 0.008991 |
| 5s/30m | 5s/15s | -2.52431 | 0.01159 |
| Naïve | 5s/15m | 2.52211 | 0.01167 |
| 5s/15m | 5s/1m | -2.48261 | 0.01304 |
| 5s/3m | 5s/1m | -2.29482 | 0.02174 |
| 5s/30s | 5s/15s | -1.91774 | 0.05514 |
| 5s/15s | 5s/15m | 1.58546 | 0.1129 |
| 5s/15s | 5s/3m | 1.53774 | 0.1241 |
| 5s/30m | 5s/3m | -1.50554 | 0.1322 |
| 5s/30m | 5s/15m | -1.10034 | 0.2712 |
| 5s/30s | 5s/3m | -0.76528 | 0.4441 |
| 5s/30s | 5s/30m | 0.60235 | 0.5469 |
| 5s/30s | 5s/15m | -0.49455 | 0.6209 |
| 5s/15m | 5s/3m | -0.37264 | 0.7094 |
| Naïve | 5s/15s | -0.18272 | 0.855 |
| Naïve | 5s/1m | 0.10088 | 0.9196 |
| 5s/15s | 5s/1m | -0.09983 | 0.9205 |
| SuppFig1-G Statistical Analysis | | | |
| Group1 | Group2 | Mann-Whitney (Z approx.) | p value |
| Naïve | 5s/5m | -15.5804 | 9.9E-55 |
| Naïve | 5s/3m | -12.9095 | 3.98E-38 |
| 5s/5m | 5s/1m | 11.8111 | 3.42E-32 |
| 5s/30s | 5s/5m | -11.5523 | 7.19E-31 |
| 5s/30m | 5s/5m | -10.6617 | 1.54E-26 |
| 5s/30s | 5s/3m | -10.5096 | 7.8E-26 |
| 5s/3m | 5s/1m | 9.8847 | 4.85E-23 |
| 5s/15s | 5s/5m | -8.4436 | 3.08E-17 |
| 5s/30m | 5s/3m | -7.3524 | 1.95E-13 |

| | | | |
|---|---|---|---|
| 5s/30s | 5s/15m | -7.3274 | 2.35E-13 |
| 5s/15s | 5s/3m | -7.1749 | 7.24E-13 |
| 5s/15m | 5s/5m | -6.8658 | 6.61E-12 |
| 5s/5m | 5s/3m | 6.7396 | 1.59E-11 |
| Naïve | 5s/15m | -5.677 | 1.37E-08 |
| 5s/30s | 5s/30m | -5.629 | 1.81E-08 |
| 5s/15m | 5s/1m | 5.1287 | 2.92E-07 |
| 5s/15s | 5s/15m | -4.6207 | 3.82E-06 |
| Naïve | 5s/30s | 4.3449 | 1.39E-05 |
| 5s/30s | 5s/1m | -4.0639 | 4.83E-05 |
| Naïve | 5s/30m | -3.3181 | 0.000906 |
| 5s/15m | 5s/3m | -3.2215 | 0.001275 |
| 5s/30m | 5s/1m | 2.9304 | 0.003385 |
| 5s/30m | 5s/15s | 2.8151 | 0.004877 |
| 5s/30m | 5s/15m | -2.6567 | 0.007891 |
| 5s/30s | 5s/15s | -1.995 | 0.04604 |
| Naïve | 5s/15s | 1.159 | 0.2464 |
| 5s/15s | 5s/1m | -1.0441 | 0.2964 |
| Naïve | 5s/1m | 0.1986 | 0.8426 |
| SuppFig2-C Statistical Analysis | | | |
| Group1 | Group2 | Mann-Whitney (Z approx.) | p value |
| Naïve | Stim. | 1.39 | 0.1645 |
| SuppFig2-D Statistical Analysis | | | |
| Group1 | Group2 | Mann-Whitney (Z approx.) | p value |
| Naïve | Stim. | 0.35 | 0.7263 |
| SuppFig2-E Statistical Analysis | | | |
| Group1 | Group2 | Mann-Whitney (Z approx.) | p value |
| Naïve | Stim. | 0.993 | 0.3205 |
| SuppFig3-A Statistical Analysis | | | |
| Group1 | Group2 | Chi-Square | p value |
| No RNAi / Dev. Stim. | Dop1R1-RNAi | 6.551 | 0.0105 |
| No RNAi / Dev. Stim. | Dop1R2-RNAi (1) | 0.868 | 0.3514 |
| No RNAi / Dev. Stim. | Dop1R2-RNAi (2) | 13.934 | 0.0002 |
| No RNAi / Dev. Stim. | Dop2R-RNAi | 13.103 | 0.0003 |
| No RNAi / Dev. Stim. | dTk99R-RNAi | 3.749 | 0.0528 |

| | | | |
|---|---|---|---|
| No RNAi / Dev. Stim. | GABA-B-R1-RNAi | 3.749 | 0.0528 |
| No RNAi / Dev. Stim. | GABA-B-R2-RNAi | 0.04 | 0.8408 |
| No RNAi / Dev. Stim. | OAMB-RNAi (1) | 51.166 | 8.49E-13 |
| No RNAi / Dev. Stim. | OAMB-RNAi (2) | 115.36 | 6.56E-27 |
| No RNAi / Dev. Stim. | Octa2R-RNAi | 7.349 | 0.0067 |
| No RNAi / Dev. Stim. | OctB1R-RNAi | 7.452 | 0.0063 |
| No RNAi / Dev. Stim. | OctB2R-RNAi | 10.903 | 0.001 |
| No RNAi / Dev. Stim. | OctB3R-RNAi | 27.617 | 1.479E-07 |
| No RNAi / Dev. Stim. | RYaR-RNAi | 16.126 | 0.0006899 |
| No RNAi / Dev. Stim. | sNPFR-RNAi | 12.176 | 0.0004841 |
| No RNAi / Dev. Stim. | TyrRI-RNAi | 6.551 | 0.0105 |
| No RNAi / Dev. Stim. | TyrRII-RNAi | 0.04 | 0.8408 |
| SuppFig3-B Statistical Analysis | | | |
| Group1 | Group2 | Steel-Dwass (Z approx.) | p value |
| OAMB-RNAi (1) | No RNAi / Dev. Stim. | 6.77118 | 2.3E-10 |
| Dop1R2-RNAi (1) | No RNAi / Dev. Stim. | 4.41525 | 0.000176 |
| OctB2R-RNAi | No RNAi / Dev. Stim. | 3.77052 | 0.002688 |
| Dop1R1-RNAi | No RNAi / Dev. Stim. | 3.48119 | 0.007886 |
| No RNAi / No Dev. Stim. | No RNAi / Dev. Stim. | 3.45797 | 0.008562 |
| OctB1R-RNAi | No RNAi / Dev. Stim. | 3.15888 | 0.02337 |
| dTk99R-RNAi | No RNAi / Dev. Stim. | -2.6897 | 0.09131 |
| OAMB-RNAi (2) | No RNAi / Dev. Stim. | 2.60867 | 0.1125 |
| Octa2R-RNAi | No RNAi / Dev. Stim. | 1.96048 | 0.4405 |
| TyrRI-RNAi | No RNAi / Dev. Stim. | 1.93484 | 0.4595 |
| TyrRII-RNAi | No RNAi / Dev. Stim. | 1.78436 | 0.5772 |
| GABA-B-R1-RNAi | No RNAi / Dev. Stim. | 1.1319 | 0.9718 |
| OctB3R-RNAi | No RNAi / Dev. Stim. | -1.13083 | 0.9721 |
| RYaR-RNAi | No RNAi / Dev. Stim. | -1.10596 | 0.9771 |
| Dop1R2-RNAi (2) | No RNAi / Dev. Stim. | -0.84937 | 0.9986 |
| Dop2R-RNAi | No RNAi / Dev. Stim. | 0.78131 | 0.9995 |
| GABA-B-R2-RNAi | No RNAi / Dev. Stim. | -0.53465 | 1 |
| sNPFR-RNAi | No RNAi / Dev. Stim. | -0.43665 | 1 |
| SuppFig3-C Statistical Analysis | | | |
| Group1 | Group2 | Steel-Dwass (Z approx.) | p value |
| OAMB-RNAi (1) | No RNAi / Dev. Stim. | -6.21639 | 9.15E-09 |
| Dop1R1-RNAi | No RNAi / Dev. Stim. | -5.72268 | 1.88E-07 |

| | | | |
|---|---|---|---|
| OctB2R-RNAi | No RNAi / Dev. Stim. | -5.47705 | 7.75E-07 |
| No RNAi / No Dev. Stim. | No RNAi / Dev. Stim. | -4.87974 | 1.89E-05 |
| Dop2R-RNAi | No RNAi / Dev. Stim. | -4.65628 | 5.68E-05 |
| OAMB-RNAi (2) | No RNAi / Dev. Stim. | -4.40964 | 0.000181 |
| Octa2R-RNAi | No RNAi / Dev. Stim. | -4.34004 | 0.000247 |
| Dop1R2-RNAi (2) | No RNAi / Dev. Stim. | -4.3306 | 0.000258 |
| Dop1R2-RNAi (1) | No RNAi / Dev. Stim. | -4.15727 | 0.000553 |
| OctB3R-RNAi | No RNAi / Dev. Stim. | -4.12664 | 0.00063 |
| GABA-B-R2-RNAi | No RNAi / Dev. Stim. | 3.64567 | 0.004327 |
| OctB1R-RNAi | No RNAi / Dev. Stim. | -3.63591 | 0.004487 |
| RYaR-RNAi | No RNAi / Dev. Stim. | 2.16819 | 0.3021 |
| dTk99R-RNAi | No RNAi / Dev. Stim. | 2.12371 | 0.3291 |
| TyrRI-RNAi | No RNAi / Dev. Stim. | -1.96325 | 0.4385 |
| sNPFR-RNAi | No RNAi / Dev. Stim. | 1.68033 | 0.6613 |
| GABA-B-R1-RNAi | No RNAi / Dev. Stim. | -1.51602 | 0.7879 |
| TyrRII-RNAi | No RNAi / Dev. Stim. | -1.50764 | 0.7939 |
| SuppFig4-A Statistical Analysis | | | |
| Group1 | Group2 | Mann-Whitney (Z approx.) | p value |
| tbh-RNAi_2>- / Dev. Stim. | tbh-RNAi_1>tdc2 / Dev. Stim. | -15.6504 | 3.3E-55 |
| tbh-RNAi_2>- / Dev. Stim. | tbh-RNAi_1>- / Dev. Stim. | -14.9047 | 3.07E-50 |
| tbh-RNAi_2>tdc2 / Dev. Stim. | tbh-RNAi_2>- / Dev. Stim. | 12.8262 | 1.17E-37 |
| tbh-RNAi_2>- / Dev. Stim. | tbh-RNAi_1>tdc2 / No Dev. Stim. | -11.9062 | 1.1E-32 |
| tbh-RNAi_2>tdc2 / Dev. Stim. | tbh-RNAi_1>tdc2 / Dev. Stim. | -11.6548 | 2.17E-31 |
| tbh-RNAi_2>tdc2 / No Dev. Stim. | tbh-RNAi_2>- / Dev. Stim. | 9.3159 | 1.21E-20 |
| tbh-RNAi_1>tdc2 / Dev. Stim. | tbh-RNAi_1>- / Dev. Stim. | 8.8655 | 7.62E-19 |
| tbh-RNAi_2>- / Dev. Stim. | tbh-RNAi_1>- / No Dev. Stim. | -8.6111 | 7.23E-18 |
| tbh-RNAi_2>tdc2 / Dev. Stim. | tbh-RNAi_1>tdc2 / No Dev. Stim. | -8.2312 | 1.85E-16 |
| tbh-RNAi_2>- / No Dev. Stim. | tbh-RNAi_1>tdc2 / Dev. Stim. | -8.0348 | 9.38E-16 |
| tbh-RNAi_2>tdc2 / No Dev. Stim. | tbh-RNAi_1>tdc2 / Dev. Stim. | -7.8141 | 5.54E-15 |

| | | | |
|---|---|---|---|
| tbh-RNAi_2>- / No Dev. Stim. | tbh-RNAi_2>- / Dev. Stim. | 7.3781 | 1.61E-13 |
| tbh-RNAi_2>- / No Dev. Stim. | tbh-RNAi_1>tdc2 / No Dev. Stim. | -6.4254 | 1.32E-10 |
| tbh-RNAi_2>tdc2 / No Dev. Stim. | tbh-RNAi_1>tdc2 / No Dev. Stim. | -5.8614 | 4.59E-09 |
| tbh-RNAi_1>tdc2 / No Dev. Stim. | tbh-RNAi_1>- / Dev. Stim. | 5.0848 | 3.68E-07 |
| tbh-RNAi_2>tdc2 / Dev. Stim. | tbh-RNAi_1>- / Dev. Stim. | -4.6646 | 3.09E-06 |
| tbh-RNAi_2>- / No Dev. Stim. | tbh-RNAi_1>- / Dev. Stim. | -3.9227 | 8.76E-05 |
| tbh-RNAi_2>- / No Dev. Stim. | tbh-RNAi_1>- / No Dev. Stim. | -3.2746 | 0.001058 |
| tbh-RNAi_2>tdc2 / Dev. Stim. | tbh-RNAi_1>- / No Dev. Stim. | -3.2127 | 0.001315 |
| tbh-RNAi_1>tdc2 / Dev. Stim. | tbh-RNAi_1>- / No Dev. Stim. | 2.9955 | 0.00274 |
| tbh-RNAi_2>tdc2 / No Dev. Stim. | tbh-RNAi_2>- / No Dev. Stim. | 2.5565 | 0.01057 |
| tbh-RNAi_2>tdc2 / No Dev. Stim. | tbh-RNAi_2>tdc2 / Dev. Stim. | 2.1526 | 0.03135 |
| tbh-RNAi_2>tdc2 / No Dev. Stim. | tbh-RNAi_1>- / No Dev. Stim. | -2.1362 | 0.03266 |
| tbh-RNAi_1>tdc2 / No Dev. Stim. | tbh-RNAi_1>- / No Dev. Stim. | 1.9785 | 0.04787 |
| tbh-RNAi_2>tdc2 / No Dev. Stim. | tbh-RNAi_1>- / Dev. Stim. | -1.6495 | 0.09905 |
| tbh-RNAi_1>tdc2 / No Dev. Stim. | tbh-RNAi_1>tdc2 / Dev. Stim. | -1.5522 | 0.1206 |
| tbh-RNAi_2>tdc2 / Dev. Stim. | tbh-RNAi_2>- / No Dev. Stim. | 1.3858 | 0.1658 |
| tbh-RNAi_1>- / No Dev. Stim. | tbh-RNAi_1>- / Dev. Stim. | 1.1529 | 0.249 |
| SuppFig4-B Statistical Analysis | | | |
| Group1 | Group2 | Mann-Whitney (Z approx.) | p value |
| tbh-RNAi_2>- / Dev. Stim. | tbh-RNAi_1>tdc2 / Dev. Stim. | 13.7706 | 3.83E-43 |
| tbh-RNAi_2>- / Dev. Stim. | tbh-RNAi_1>tdc2 / No Dev. Stim. | 12.3092 | 8.08E-35 |
| tbh-RNAi_2>tdc2 / Dev. Stim. | tbh-RNAi_1>tdc2 / No Dev. Stim. | 11.0635 | 1.89E-28 |
| tbh-RNAi_2>tdc2 / No Dev. Stim. | tbh-RNAi_2>- / Dev. Stim. | -10.8111 | 3.05E-27 |

| | | | |
|---|---|---|---|
| tbh-RNAi_1>tdc2 / No Dev. Stim. | tbh-RNAi_1>- / Dev. Stim. | -10.2397 | 1.32E-24 |
| tbh-RNAi_2>- / Dev. Stim. | tbh-RNAi_1>- / Dev. Stim. | 10.2067 | 1.85E-24 |
| tbh-RNAi_2>- / Dev. Stim. | tbh-RNAi_1>- / No Dev. Stim. | 10.0062 | 1.43E-23 |
| tbh-RNAi_2>- / No Dev. Stim. | tbh-RNAi_2>- / Dev. Stim. | -9.3501 | 8.76E-21 |
| tbh-RNAi_2>tdc2 / Dev. Stim. | tbh-RNAi_1>- / No Dev. Stim. | 9.2063 | 3.38E-20 |
| tbh-RNAi_1>tdc2 / No Dev. Stim. | tbh-RNAi_1>tdc2 / Dev. Stim. | -8.9635 | 3.14E-19 |
| tbh-RNAi_2>tdc2 / Dev. Stim. | tbh-RNAi_1>tdc2 / Dev. Stim. | 8.8678 | 7.46E-19 |
| tbh-RNAi_1>- / No Dev. Stim. | tbh-RNAi_1>- / Dev. Stim. | -8.4605 | 2.66E-17 |
| tbh-RNAi_1>tdc2 / Dev. Stim. | tbh-RNAi_1>- / No Dev. Stim. | 7.8068 | 5.86E-15 |
| tbh-RNAi_2>tdc2 / No Dev. Stim. | tbh-RNAi_2>tdc2 / Dev. Stim. | -7.3893 | 1.48E-13 |
| tbh-RNAi_2>- / No Dev. Stim. | tbh-RNAi_1>tdc2 / No Dev. Stim. | 6.8782 | 6.06E-12 |
| tbh-RNAi_2>tdc2 / No Dev. Stim. | tbh-RNAi_1>tdc2 / No Dev. Stim. | 6.75 | 1.48E-11 |
| tbh-RNAi_2>- / No Dev. Stim. | tbh-RNAi_1>- / No Dev. Stim. | 6.6336 | 3.28E-11 |
| tbh-RNAi_2>tdc2 / No Dev. Stim. | tbh-RNAi_1>- / No Dev. Stim. | 6.3796 | 1.78E-10 |
| tbh-RNAi_2>tdc2 / Dev. Stim. | tbh-RNAi_2>- / Dev. Stim. | -6.1407 | 8.22E-10 |
| tbh-RNAi_2>tdc2 / Dev. Stim. | tbh-RNAi_2>- / No Dev. Stim. | 5.9568 | 2.57E-09 |
| tbh-RNAi_1>tdc2 / Dev. Stim. | tbh-RNAi_1>- / Dev. Stim. | -5.1479 | 2.63E-07 |
| tbh-RNAi_2>tdc2 / Dev. Stim. | tbh-RNAi_1>- / Dev. Stim. | 4.345 | 1.39E-05 |
| tbh-RNAi_2>tdc2 / No Dev. Stim. | tbh-RNAi_1>- / Dev. Stim. | -4.1968 | 2.71E-05 |
| tbh-RNAi_2>- / No Dev. Stim. | tbh-RNAi_1>- / Dev. Stim. | -3.0713 | 0.002131 |
| tbh-RNAi_2>tdc2 / No Dev. Stim. | tbh-RNAi_2>- / No Dev. Stim. | -0.8008 | 0.4233 |
| tbh-RNAi_2>tdc2 / No Dev. Stim. | tbh-RNAi_1>tdc2 / Dev. Stim. | -0.7264 | 0.4676 |
| tbh-RNAi_1>tdc2 / No Dev. Stim. | tbh-RNAi_1>- / No Dev. Stim. | 0.6361 | 0.5247 |

| | | | |
|---|---|---|---|
| tbh-RNAi_2>- / No Dev. Stim. | tbh-RNAi_1>tdc2 / Dev. Stim. | 0.2063 | 0.8366 |
| SuppFig4-C Statistical Analysis | | | |
| Group1 | Group2 | Mann-Whitney (Z approx.) | p value |
| tdc2-GAL4 / Dev. Stim. | attp2 / No Dev. Stim. | -10.623 | 2.33E-26 |
| tdc2-GAL4 / Dev. Stim. | attp2 / Dev. Stim. | -7.8858 | 3.13E-15 |
| attp2 / No Dev. Stim. | attp2 / Dev. Stim. | 7.1702 | 7.49E-13 |
| tdc2-GAL4 / No Dev. Stim. | attp2 / No Dev. Stim. | -6.6541 | 2.85E-11 |
| tdc2-GAL4 / No Dev. Stim. | tdc2-GAL4 / Dev. Stim. | 3.7126 | 0.000205 |
| tdc2-GAL4 / No Dev. Stim. | attp2 / Dev. Stim. | -2.8086 | 0.004975 |
| SuppFig4-D Statistical Analysis | | | |
| Group1 | Group2 | Mann-Whitney (Z approx.) | p value |
| tdc2-GAL4 / Dev. Stim. | attp2 / No Dev. Stim. | 14.5498 | 5.86E-48 |
| tdc2-GAL4 / No Dev. Stim. | tdc2-GAL4 / Dev. Stim. | -14.5297 | 7.86E-48 |
| attp2 / No Dev. Stim. | attp2 / Dev. Stim. | -13.4172 | 4.8E-41 |
| tdc2-GAL4 / No Dev. Stim. | attp2 / Dev. Stim. | -12.1437 | 6.2E-34 |
| tdc2-GAL4 / Dev. Stim. | attp2 / Dev. Stim. | 6.6036 | 4.01E-11 |
| tdc2-GAL4 / No Dev. Stim. | attp2 / No Dev. Stim. | 5.5707 | 2.54E-08 |
